# Predmoter - Cross-species prediction of plant promoter and enhancer regions

**DOI:** 10.1101/2023.11.03.565452

**Authors:** Felicitas Kindel, Sebastian Triesch, Urte Schlüter, Laura Alexandra Randarevitch, Vanessa Reichel-Deland, Andreas P.M. Weber, Alisandra K. Denton

**Author notes:** Corresponding author: Alisandra Denton. **Email addresses:**.

## Abstract

**Motivation:** The identification of *cis*-regulatory elements (CREs) is crucial for the analysis of gene regulatory networks in plants. Several next generation sequencing (NGS)-based methods were developed to identify CREs. However, these methods can be time-consuming and costly. They also involve creating sequencing libraries for the entire genome. Since many research efforts only focus on specific genomic loci, this presents a considerable expenditure. Computational prediction of the outputs of specialized NGS methods to analyze CREs, like Assay for Transposase Accessible Chromatin using sequencing (ATAC-seq), would significantly cut costs and time investment. Yet, no such method is available to date.

**Results:** We present Predmoter, a deep neural network able to predict base-wise ATAC-seq and histone Chromatin immunoprecipitation DNA-sequencing (ChIP-seq) read coverage for plant genomes. Predmoter uses only the DNA sequence as input. We evaluated our model on two plant genomes, the genome of the dicot *Arabidopsis thaliana* and of the monocot *Oryza sativa*. We trained our models on 10 species with publicly available ATAC-seq data and 15 species with ChIP-seq data. Our best models showed accurate predictions in peak positions and the overall pattern of peaks for ATAC- and Histone H3 trimethylated at lysine 4 (H3K4me3) ChIP-seq. Annotating putatively accessible chromatin regions provides valuable input for the identification of CREs. In conjunction with other *in silico* data, such as predicted binding affinities for transcription factors (TFs), this can significantly narrow down the search space to a manageable number of experimentally verifiable DNA-protein interaction pairs.

**Availability and Implementation:** The source code for Predmoter is available at: https://github.com/weberlab-hhu/Predmoter along with documentation for installation and usage. Predmoter uses a single-command inference, Predmoter.py, for both training and prediction. Predmoter takes a fasta file as input and outputs an h5 file and optionally bigWig and bedGraph files.

**Highlight:** Predmoter will help identifying CREs and so gaining further insight into gene regulatory networks in plants.

## 1. Introduction

Despite large genomic and epigenomic studies being published in all fields of biology, the identification of *cis*-regulatory sequences and their influence on gene regulation is still a major challenge. The discovery of new *cis*-regulatory elements (CREs) can reveal targets for genetic engineering and breeding supporting optimization of plant growth as well as stress and pathogen resistance.

Two important locations of CREs are promoters and enhancers. Promoters are historically defined to serve transcription initiation (Ippen et al. 1968, Epstein and Beckwith 1968, Jacob, Ullman and Monod 1964). The core promoter is a region of 50 to 100 base pairs (bp) upstream from the transcription start site (TSS) (Struhl 1995, Dynan and Tjian 1985). We refer here to promoter as the assembly of individual transcription factor (TF) binding sites, i.e., CREs, upstream of a gene that entirely or partially drive local transcription initiation. This region contains at least the core promoter. On the other hand, enhancers can increase transcription levels from a given promoter. They were found to act in either orientation and at many positions. The first discovered enhancer sequence was found in *Escherichia coli*, and it could act up to 1400 bp upstream or 3300 bp downstream from the TSS (Banerji, Rusconi and Schaffner 1981). An example distal enhancer in plants is acting 140 kbp upstream of the *bx1* gene in *Zea mays* (Zheng et al. 2015). Whereas the core promoter mostly coordinates expression of the adjesent gene, enhancers can regulate gene expression of multiple genes.

The binary classification of promoters and enhancers has since been challenged. Promoters with high enhancer strengths (Dao et al. 2017, Diao et al. 2017, Engreitz et al. 2016) and active enhancers driving local transcription initiation at their boundaries (Andersson et al. 2014, de Santa et al. 2010, Kim et al. 2010) have been reported. Promoters and enhancers usually are both found in accessible chromatin regions (ACRs), where the DNA is accessible to TFs (Cockerill 2011, Gross and Garrard 1988, Song et al. 2011). Both promoter and enhancer regions are marked by different histone modifications. Histone H3 trimethylated at lysine 4 (H3K4me3) is primarily present at active genes, while H3K4me2 occurs at both inactive and active euchromatic genes (Santos-Rosa et al. 2002). Both can be detected in the core promoter and the coding region of genes. Enhancers are instead marked by H3K4me1 (Heintzman et al. 2009). Active enhancers are additionally marked by an acetylation of H3K27 (H3K27ac) (Rada-Iglesias et al. 2010, Creyghton et al. 2010). Poised or inactive enhancers are in contrast marked by the absence of H3K27ac, instead showing an enrichment of H3K27 trimethylation (H3K27me3) (Rada-Iglesias et al. 2010, Creyghton et al. 2010). However, H3K4me1 was found to not commonly be associated with distal ACRs in plants (Lu et al. 2019).

Assay for Transposase Accessible Chromatin using sequencing (ATAC-seq) is a common method to identify ACRs (Buenrostro et al. 2013). It is faster and more sensitive than previous methods like DNase I hypersensitive sites sequencing (DNase-seq) (Crawford et al. 2006) or formaldehyde-assisted isolation of regulatory elements (FAIRE-seq) (Giresi et al. 2007). ATAC-seq uses hyperactive mutant Tn5-transposase, which cuts the DNA primarily in ACRs and ligates adapters to the cut DNA fragment (Buenrostro et al. 2013). The resulting fragments are amplified by PCR creating a sequencing library. In contrast to ATAC-seq, which outputs ACRs, chromatin immunoprecipitation DNA-sequencing (ChIP-seq) (Robertson et al. 2007, Johnson et al. 2007, Kim et al. 2004) is used to investigate how proteins that interact with the DNA regions of interest regulate target gene expression. Proteins attached to the DNA are crosslinked with the DNA, the DNA is sheared, the proteins are immunoprecipitated and unlinked, so the DNA can be amplified and sequenced (Robertson et al. 2007, Johnson et al. 2007, Kim et al. 2004). Depending on the assay, either TF or histone antibodies are used in immunoprecipitation. Promoter as well as enhancer specific histone modifications can be identified using ChIP-seq.

Deep learning (DL) is a part of machine learning using artificial neural networks (NNs) that have multiple hidden layers creating a deep neural network (DNN) architecture (Schulz and Behnke 2012).

*In silico* identification of promoter and enhancer sequences using DL was attempted in several studies. Most studies predicted promoters as a sequence stretch around the TSS (Umarov and Solovyev 2017, Shahmuradov, Umarov and Solovyev 2017, Umarov et al. 2019, Oubounyt et al. 2019, Shujaat et al. 2021, Wang et al. 2022). The networks in these studies performed a fundamentally different predictive task than actual promoter sequence prediction. Other approaches of predicting regulatory factor binding activity (Hiranuma, Lundberg and Lee 2017) or predicting enhancer regions (Thibodeau et al. 2018) utilized ATAC-seq data in conjunction with DNA sequence information. However, these networks only utilize ATAC-seq data from human samples. Furthermore, the Enformer DNN can predict gene expression and chromatin states, represented as multiple genomic coverage tracks like H3K27ac coverage, in humans and mice from DNA sequences (Avsec et al. 2021). Plant research keeps lagging behind research in mammalian species in this field and a DNN focused on predicting plant CREs would be a first step to alleviate this underrepresentation. Moreover, generating ATAC- and ChIP-seq libraries is costly and time consuming and a DNN predicting plant ATAC- and ChIP-seq read coverage directly from the genomic DNA sequence would circumvent these constraints. To date, no such model has been reported.

Here we present Predmoter, a tool used for cross-species base-wise prediction of plant ATAC- and/or H3K4me3 ChIP-seq read coverage, using the genomic DNA sequence as input. We utilized publicly available ATAC- and ChIP-seq data to infer plant promoter and enhancer regions. We trained our model on ATAC-seq data from 10 different plant species and ChIP-seq data from 15 plant species.

## 2. System and methods

### 2.1 Data

The entire dataset consisted of 22 plant genomes, publicly available ATAC-seq reads from 14 plant species and publicly available ChIP-seq (H3K4me3) reads from 19 plant species (see Table 1 and Supplementary Table S2). A wide variety of tissues and treatments were used in these ATAC- and ChIP-seq experiments which are listed in Supplementary Table S3.

**Table 1:**
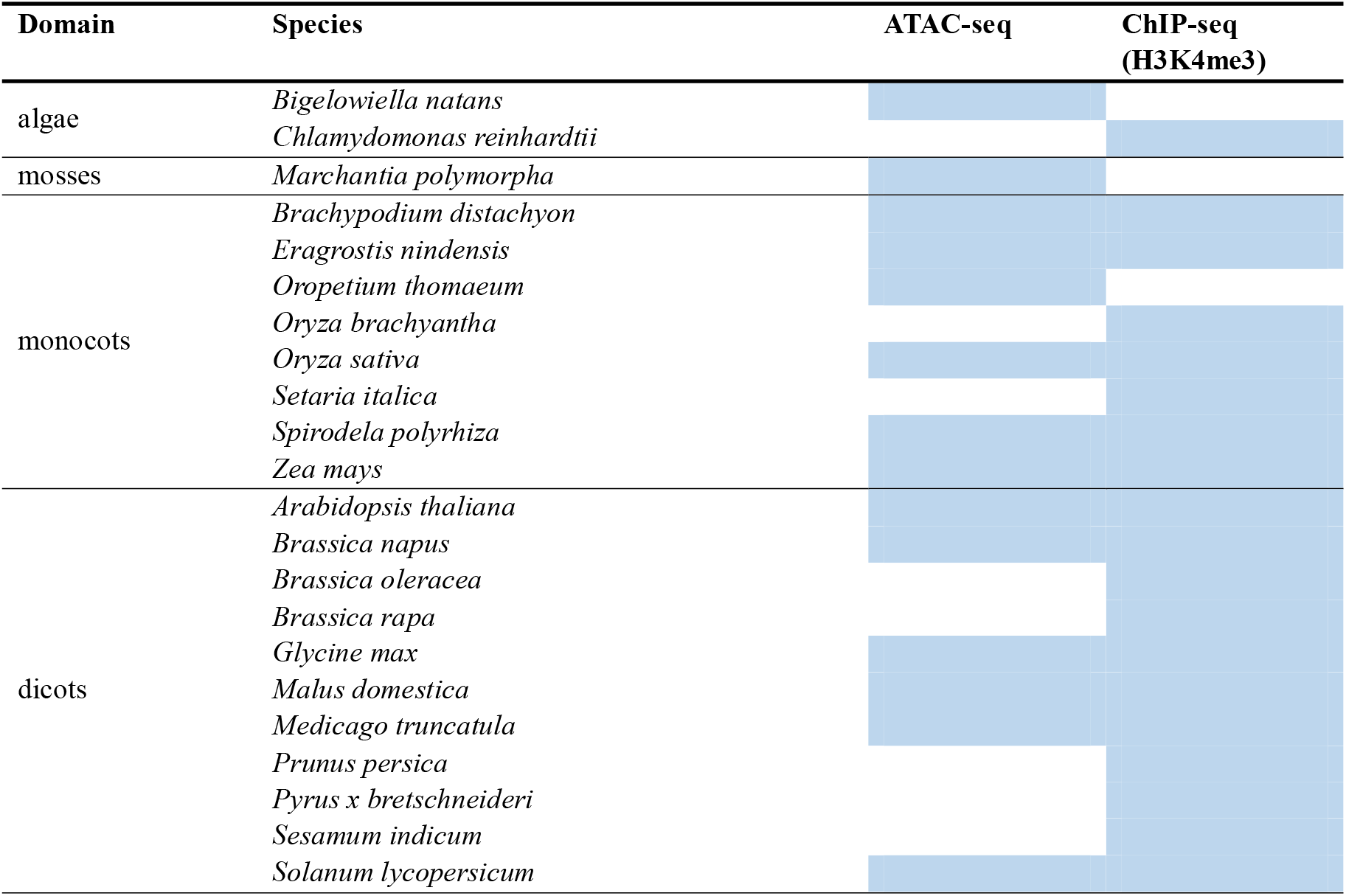
Plant genomes and available datasets. The species used in the development of Predmoter are separated into the four domains algae, mosses, monocots, and dicots. The availability and usage of the species dataset for ATAC- or ChIP-seq is indicated by a blue box.

The NGS data was downloaded from the sequence read archive (SRA) using the SRA-Toolkit 3.0.0 (https://github.com/ncbi/sra-tools/wiki/01.-Downloading-SRA-Toolkit). The reads were trimmed with Trimmomatic 0.36 (Bolger, Lohse and Usadel 2014) and quality controlled using FastQC 0.11.9 (Andrews 2010) and MultiQC (Ewels et al. 2016). If the reads passed quality control, they were mapped to the reference genome using BWA 2.1 (Md et al. 2019). Conversion to bam files was performed using SamTools 1.6 (Danecek et al. 2021). The Picard Toolkit (Broad Institute 2019) was used to mark duplicates. The duplicates, unmapped reads, non-primary alignments and reads not passing platform quality checks were removed with SamTools. Plots for quality control were generated using deepTools 3.5.3 (Ramírez et al. 2016) and the necessary genome annotations were generated using Helixer v.0.3.1 (Stiehler et al. 2021, Holst et al. 2023). A detailed data preprocessing documentation is available at: https://github.com/weberlab-hhu/Predmoter/blob/main/docs/data_preprocessing.md. The plant genome fasta files and final NGS data bam files were converted to h5 files using Helixer (Stiehler et al. 2021, Holst et al. 2023). The ATAC-seq reads were shifted +4 bp on the positive strand and -5 bp on the negative strand to adjust the read start sites to represent the center of the transposon binding site (Buenrostro et al. 2013). A detailed documentation of the h5 file creation and architecture is available at: https://github.com/weberlab-hhu/Predmoter/blob/main/docs/h5_files.md. Peak calling on predictions and the experimental data was performed with MACS3 (Zhang et al. 2008). For ATAC-seq “bdgpeakcall” was used with a minimum read length of 85, maximum gap size of 35 and a cutoff of 10 for *A. thaliana* and with a minimum read length of 60, maximum gap size of 70 and a cutoff of 10 for *O. sativa*. For ChIP-seq “bdgbroadcall” was used with a minimum read length of 180, maximum gap size of 60, a maximum linking gap of 720, a cutoff of 120 and a linking cutoff of 20 for *A. thaliana*. For *O. sativa* the minimum read length was 145, the maximum gap size 95, the maximum linking gap 580, the cutoff 120 and the linking cutoff 15. A detailed explanation of these parameter choices can be found in Supplementary Section 1.7.

Ensuring a diverse range of species in the training set, while simultaneously reserving enough data for validation and testing to effectively evaluate the models’ generalization ability, proved difficult. At the time of development, the amount of high-quality, publicly available ATAC-seq data was low. Around 60 % of the plant ATAC-seq data on SRA available up until July 2023 needed to be discarded after the final quality control. The ATAC-seq coverage enrichment +/- 1.5 kbp around the TSS either did not show the correct or any peak around or upstream of the TSS, or the average peak read coverage was less than half of the background coverage. This left the ATAC-seq data of the 14 plant species used in this study. The quality control for ChIP-seq data was performed using the same criteria. The low availability of high-quality data turned out to be a major hindrance in providing the network with an appropriate amount of data to train on. Data of two species, *A. thaliana* and *O. sativa*, was set aside as a hold-out test set. In doing so, both a dicot and a monocot species with available ATAC- and ChIP-seq data could be used for final evaluation. The same was done with the two validation species, the dicot *Medicago truncatula* and the monocot *Spirodela polyrhiza* (see Supplementary Table S4). Cross-species validation instead of an in-species split for the validation and training data was deemed closer to the real-world use case of predicting ATAC- and ChIP-seq data for an entire species.

### 2.2 Architecture and training

The model architectures were implemented using Pytorch Lightning (Falcon 2019) on top of PyTorch (Paszke et al. 2019). The model used supervised learning, a method that connects an input to an output based on example input-output pairs (Russell and Norvig 2016).

The input for the model was a genomic DNA sequence. The nucleotides were encoded into four-dimensional vectors (see Supplementary Table S1). The DNA sequence of a given plant species was cut into subsequences of 21384 bp. As few chromosomes, scaffolds or contigs were divisible by this number, sequence ends as well as short sequences were padded with the vector [0., 0., 0., 0.]. Padded base pairs were masked during training. If a subsequence only contained N bases, here referred to as “gap subsequence”, it was filtered out. Both strands, plus and minus, were used. Since the ATAC- and ChIP-seq data was PCR amplified and as such it was not possible to determine from which strand a read originated, the coverage information was always added to both strands. The model’s predictions for either ATAC-seq, ChIP-seq or both were compared to the experimental read coverage. The target data was represented per sample of experimental data. These were averaged beforehand, resulting in one coverage track per NGS dataset and plant species.

Three main model architectures were examined on their performance. The first architecture consisted of convolutional layers followed by transposed convolutional layers for deconvolution (LeCun and Bengio 1995, LeCun et al. 1989). The deconvolution was added to output base-wise predictions. We refer here to this architecture as U-Net. Our second approach was a hybrid network. A block of long short-term memory layers (LSTM) (Hochreiter and Schmidhuber 1997) was placed in between a convolutional layer block and a transposed convolutional layer block. The final approach was called bi-hybrid. Its architecture matched the hybrid architecture, except that the LSTM layers were replaced with bidirectional LSTM layers (BiLSTM) (Hochreiter and Schmidhuber 1997, Schuster and Paliwal 1997). Each convolutional and transposed convolutional layer was followed in all three approaches by the ReLU activation function (Glorot, Bordes and Bengio 2011). Additional augmentations to the bi- hybrid network included adding batch normalization after each convolutional and transposed convolutional layer and adding a dropout layer after each BiLSTM layer except the last (Fig. 1). The Adam algorithm was used as an optimization method (Kingma and Ba 2014).

**Figure 1:**
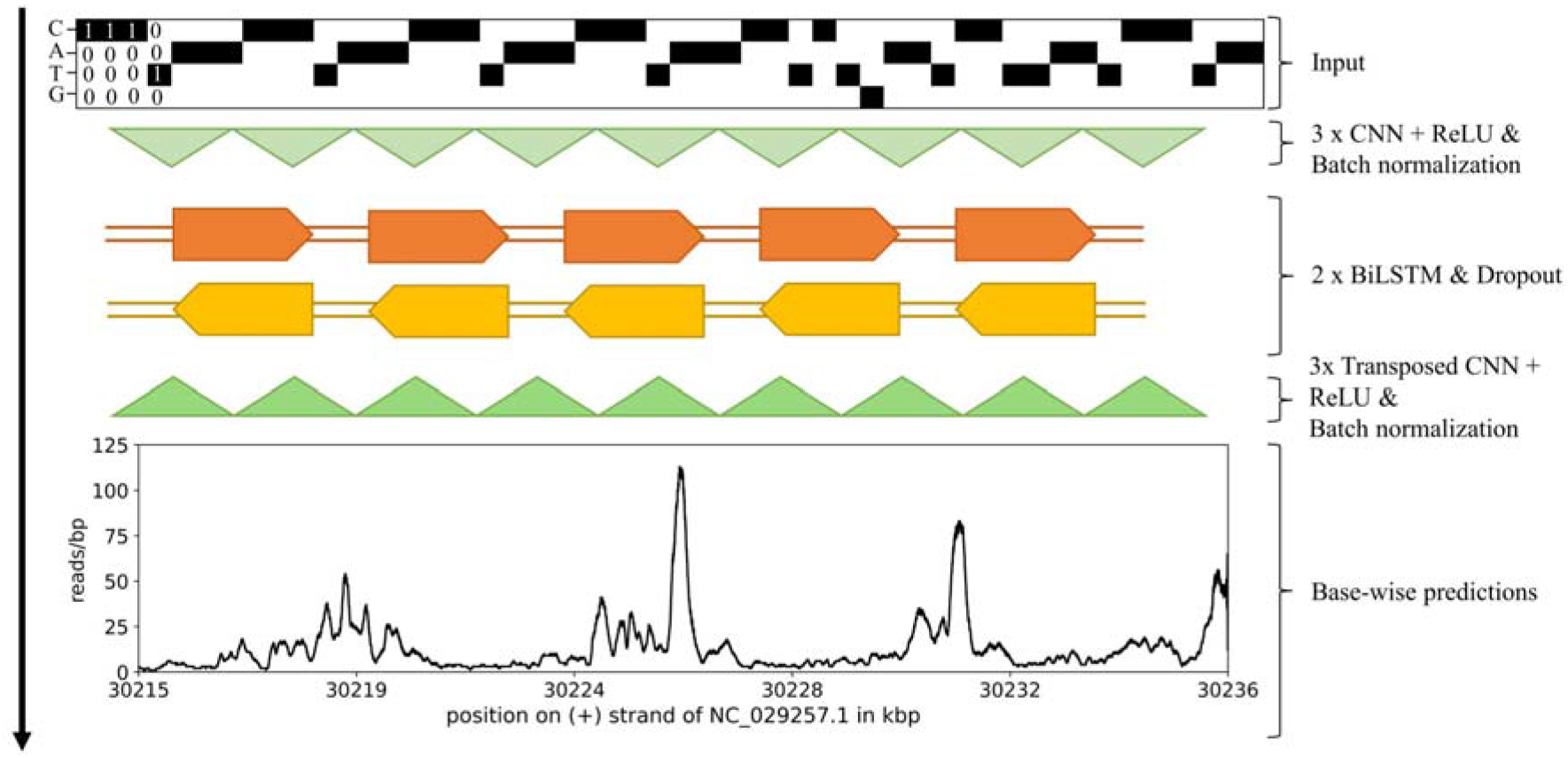
Predmoter architecture and prediction process. The bi-hybrid architecture with batch normalization and dropout is schematically depicted. Not to scale. Hyperparameters are examples and can vary. The base-wise predictions are from an example subsequence from *O. sativa*.

Since convolutional layers reduce the sequence length, the transposed convolutional layers were used to regain the initial sequence length to predict at full base-wise resolution. To ensure that the new sequence length resulting from a convolution was correct, custom padding formulas were used (Supplementary Section S1.2). As an additional improvement, unplaced scaffolds and non-nuclear sequences were flagged and optionally filtered out, so they were excluded from training. Sequences from genome assemblies just containing unplaced scaffolds or contigs were not flagged (Supplementary Section S1.5).

### 2.3 Metrics

Two metrics were used to evaluate model performance, the Poisson loss and the Pearson correlation coefficient (Pearson’s r).

The most prominent peak caller for ChIP-seq data, MACS (Zhang et al. 2008), which was also frequently used for ATAC-seq data (Thibodeau et al. 2018, Hiranuma, Lundberg and Lee 2017, Hentges et al. 2022), assumes that the ChIP-seq coverage data is Poisson distributed. Therefore, PyTorch’s Poisson negative log likelihood loss function (Poisson loss) was used as the loss function for all models (Equation 1). This version of the Poisson loss caused the network to output logarithmic predictions. The desired, actual predictions were thus the exponential of the network’s output. The exponential distribution only consists of positive real numbers like the ATAC- and ChIP-seq read coverage.

Equation 1: Calculation of the Poisson negative log likelihood loss

The individual samples of the predictions (x) and the targets (y) are indexed with *i*,. The sample size is denoted with *n* (https://pytorch.org/docs/stable/generated/torch.nn.PoissonNLLLoss.html).

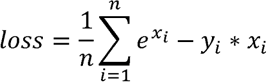

To measure the “accuracy” of the model’s predictions, i.e., translating the Poisson loss into a human-readable number, the Pearson’s r was chosen (Equation 2), measuring the linear correlation between two variables. A value of 1 represents a perfect positive linear relationship, so Predmoter’s predictions and the experimental NGS coverage data would be identical. A value of 0 means no linear relationship between the predictions and targets. Finally, a value of -1 represents a perfect negative linear relationship.

Equation 2: Calculation of the Pearson correlation coefficient (Pearson’s r)

The sample size is denoted with n, the individual samples of the predictions (x) and the targets (y) are indexed with,. The additional epsilon () equals 1e-8 and is used to avoid a division by zero.

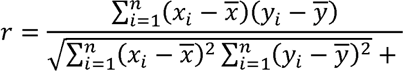

The F_1_ was used to compare predicted peaks to experimental peaks for both test species (Equation 3). A F_1_ score of 1 indicates that the predicted peaks are at the same position as the experimental peaks. The lowest score possible is 0. The F_1_ was calculated base-wise. Called peaks were denoted with 1, all other base pairs with 0. A confusion matrix containing the sum of True Positives (TP), False Positives (FP) and False Negatives (FN) for the two classes, peak and no peak, was computed for each strand.

The two matrices were then summed up. Flagged sequences were excluded from the calculations (see Supplementary Section S1.5).

Equation 3: Calculation of precision, recall and F1.

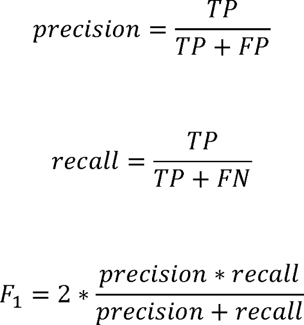

### 2.4 Usage

Inference can be done using a fasta or a h5 file containing a genome sequence as input. The h5 files are created with the *ab initio* gene caller Helixer, also developed by our institute (https://github.com/weberlab-hhu/Helixer). Helixer is implemented into Predmoter to allow direct conversion from fasta to h5 file format. The subsequence length of the h5 file can be higher or shorter than the default 21384 bp, if it is evenly divisible by the stride (step) to the power of the number of convolutional layers of the model. The final output of Predmoter is an h5 file containing information about the species and the sequence ID per subsequence, the position of the subsequence on the chromosome/sequence and the base-wise predictions of the NGS dataset or datasets used during training. The order of these datasets and the name of the model checkpoint used to generate the predictions are also saved to the h5 file. Additionally, the predictions can be converted to bedGraph or bigWig files, either via convert2coverage.py or by setting the “-of” parameter of Predmoter to either “bg” or “bw” when predicting. The strand, either the average of both, plus or minus, can be chosen in convert2coverage.py. For direct conversion after predicting, the average of both strands is chosen by default. Padded base pairs are set to the filler value of -1 in the h5 output file and excluded in the bedGraph and bigWig files. Predmoter also creates several log files (see Supplementary Section S1.4). Predmoter was built with the intention to be able to learn multiple different NGS datasets simultaneously if they follow a Poisson distribution.

### 2.5 Model releases

The exact species selection can be found in Supplementary Table S4 and the parameters in Supplementary Table S5. A detailed explanation of all available parameters can be found at: https://github.com/weberlab-hhu/Predmoter/blob/main/docs/Predmoter_options.md. The models are available at: https://github.com/weberlab-hhu/predmoter_models. The standard model naming convention of Predmoter is predmoter_<VERSION>_epoch= <EPOCH>_<CHECKPOINT_QUANTITY>.ckpt. An example of this convention is “predmoter_v0.3.2_epoch=49_avg_val_accuracy=0.5323.ckpt”. The naming convention of the released models is <MODEL_NAME>_predmoter_<PREDMOTER_VERSION>.ckpt, e.g. “BiHybrid_04_predmoter_v0.3.2.ckpt”. The released models were the models with the highest validation Pearson’s correlation. They were used for testing and in some cases inference.

## 3. Results

Nine different models were evaluated (Tab. 2). The best model of each architecture and dataset combination was used to develop the next combination test. The model reaching the highest Pearson’s correlation for the validation set was deemed the best model. Pre-tests showed that including gap subsequences, subsequences of 21384 bp only containing Ns, led to a considerably lower Pearson’s correlation. The proportion of gap subsequences in the total data was 0.6 %. Normalizing the NGS coverage data through a general approach of subtracting the average coverage from the dataset and using a ReLU transformation (Glorot, Bordes and Bengio 2011) showed notably worse results during previous attempts. The approach of normalizing via an input sample was not feasible due to the considerable lack of available ATAC-seq input samples accompanying the experiments. Therefore, the target data was not adjusted towards its sequencing depth.

**Table 2:** Model architecture and dataset explanation (short). All models excluded gap subsequences, subsequences of 21384 bp only containing Ns. For more details on species selection and exact model parameters see Supplementary Tables S4 and S5.

| Model name | Dataset | Architecture |
| --- | --- | --- |
| U-Net | ATAC-seq | U-Net |
| Hybrid | ATAC-seq | U-Net + LSTM |
| BiHybrid | ATAC-seq | U-Net + BiLSTM |
| BiHybrid_02 | ATAC-seq | U-Net + BiLSTM + batch normalization |
| BiHybrid_03.1 | ATAC-seq | U-Net + BiLSTM + batch normalization + dropout of 0.3 |
| BiHybrid_03.2 | ATAC-seq | U-Net + BiLSTM + batch normalization + dropout of 0.5 |
| BiHybrid_04 | ATAC-seq, filtered flagged sequences* | U-Net + BiLSTM + batch normalization + dropout of 0.3 |
| BiHybrid_05 | ChIP-seq (H3K4me3), filtered flagged sequences* | U-Net + BiLSTM + batch normalization + dropout of 0.3 |
| Combined | ATAC-seq, ChIP-seq (H3K4me3), filtered flagged sequences* | U-Net + BiLSTM + batch normalization + dropout of 0.3 |

\*excluded subsequences of unplaced scaffolds and non-nuclear sequences during training and testing

A comparison of the first three setups showed that the best base architecture was the BiLSTM layers placed in between a block of convolutional layers and a block of transposed convolutional layers, called “bi-hybrid” in Predmoter (Fig. 2). The architecture used three convolutional, three transposed convolutional and two BiLSTM layers. This setup outperformed the U-Net architecture, which was missing the LSTM layers in the middle, as well as the hybrid architecture that utilized two one-directional LSTM layers. The U-Net performed worst out of all examined models. The model setup BiHybrid_02 added batch normalization after each convolutional and transposed convolutional layer. These additional six layers improved the results further. Introducing a dropout layer with a dropout probability of 30 % between the two LSTM layers, model architecture BiHybrid_03.1, showed modest improvements. In contrast, the architecture BiHybrid_03.2 with a dropout probability of 50 % did not improve the model. Filtering flagged sequences, meaning unplaced scaffolds and non-nuclear sequences, in the assembly where possible, was introduced for model ByHybrid_04. Filtering improved the test metrics slightly compared to BiHybrid_03.1. For this comparison the flagged sequences were also once excluded during testing, but not training of BiHybrid_03.1. This approach was chosen, because non-nuclear sequences and unplaced scaffolds were observed to cause high amounts of background noise in the ATAC-seq experimental data during data quality control in some cases. The unplaced scaffolds and non-nuclear sequences reached 7.1 % of all genome assemblies used not counting assemblies on scaffold or contig level, as they were not filtered.

**Figure 2:**
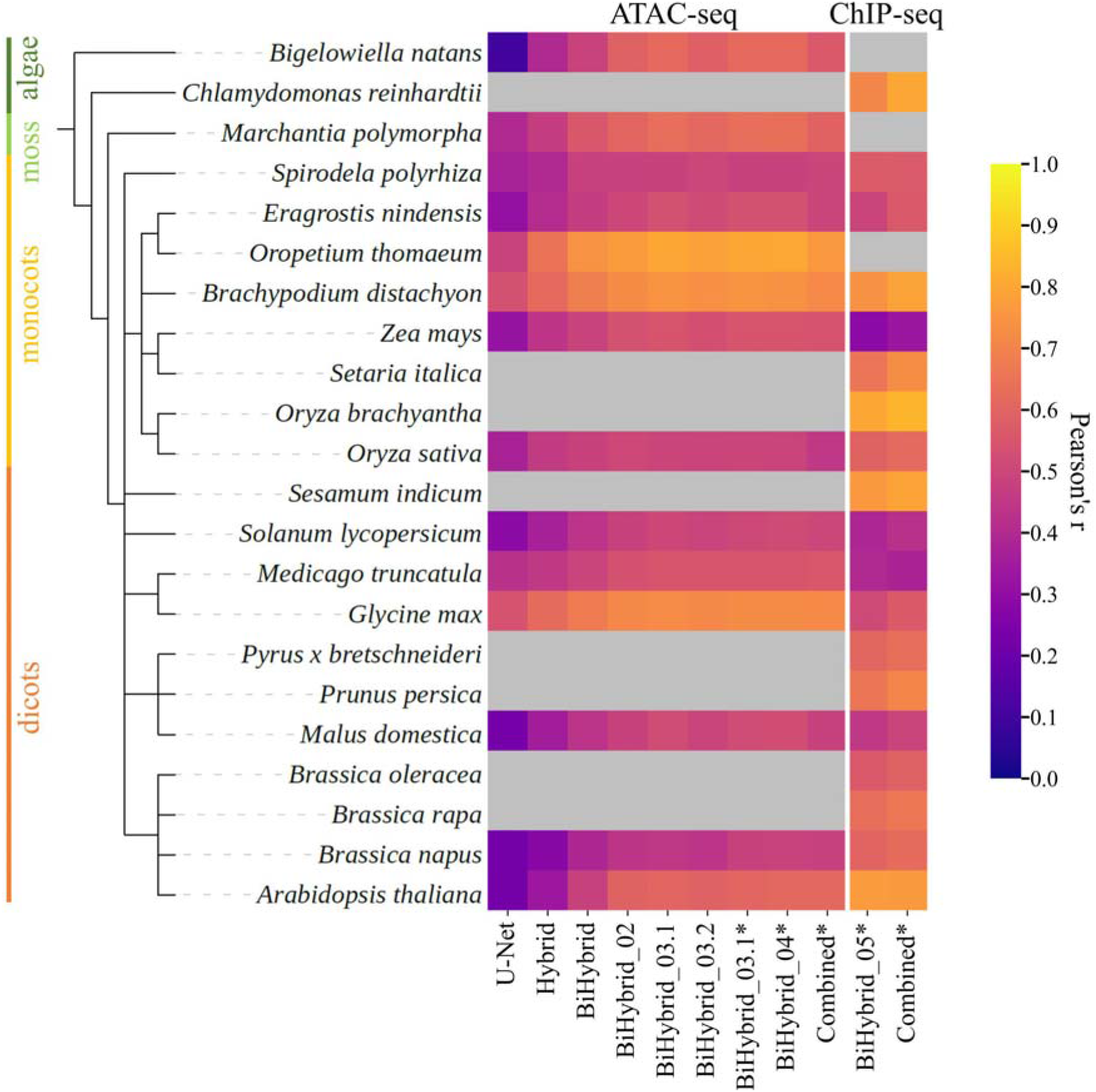
Performance of the best models per model setup across all species. The performance is measured via the Pearson correlation coefficient by comparing the experimental data (target) with the model’s prediction. Gap subsequences were excluded during testing. Results marked with * also excluded flagged subsequences. The left block shows the results for ATAC-seq, the right one for ChIP- seq (H3K4me3). Grey boxes are used when there was no available high-quality experimental data for the given NGS dataset and species to compare predictions to. Tabular results are listed in Supplementary Tables S7 and S8.

This final stage of the models’ architecture and development was then used to train on ChIP-seq (H3K4me3) data instead of ATAC-seq data, denoted as model BiHybrid_05. A final combined model was trained using the setup of BiHybrid_04 and BiHybrid_05, but training on ATAC- and ChIP-seq data simultaneously. For the ChIP-seq data, noise originating from non-nuclear sequences and unplaced scaffolds was not observed. The flagged data therefore would have been for the most part another set of the “negative” data with no associated ChIP-seq peaks. As the ATAC- and ChIP-seq data cannot be filtered independently in Predmoter’s implementation, filtering of flagged sequences was used for both the BiHybrid_05 and the combined model to ensure comparability. The combined model performed better on the ChIP-seq data than the ChIP-seq model BiHybrid_05, but worse for the ATAC-seq data than the previous best ATAC-seq model BiHybrid_04. There was no bias towards dicots. Dicots made up 50.8 % of the training set after removing gap and flagged subsequences, while monocots made up 43.9 %. Algae and mosses only made up 2.5 % and 2.8 % of the training set respectively.

Coverage enrichment +/- 3 kbp around the TSS of the ATAC- and ChIP-seq predictions and experimental data from *A. thaliana* and *O. sativa* showed that the predicted peaks had the same pattern and were at the same location as the ones from the experimental data (Fig. 3). For all models the read coverage was on average predicted to be up to 50 reads per bp lower for the background coverage (Fig. 3b) and up to 175 reads per bp lower in peak regions (Fig. 3b) than suggested by the available experimental data. The amplitude of the predicted ATAC-seq data around the TSS was 16 % higher for the combined model than for BiHybrid_03.1 or BiHybrid_04 (Fig. 3c). The ATAC-seq peak predictions for *A. thalina* appeared to be notably below the experimental data in terms of coverage (Fig. 3a). However, the pattern and position if not the width of the predicted peaks was like the experimental data. For the ChIP-seq data the amplitudes from the two *A. thaliana* predictions were similar (Fig. 3b). Meanwhile, for *O. sativa* the predictions of the BiHybrid_05 model showed a on average 7 reads per bp higher amplitude than the ones of the combined model (Fig. 3d). All predictions for *A. thaliana* showed a lower read coverage per bp than the ones of *O. sativa*. The ATAC-seq peak predictions also all had a lower amplitude than the ChIP-seq predictions. This mimicked the lower amplitudes of the experimental ATAC-seq data in comparison to the experimental ChIP-seq data.

**Figure 3:**
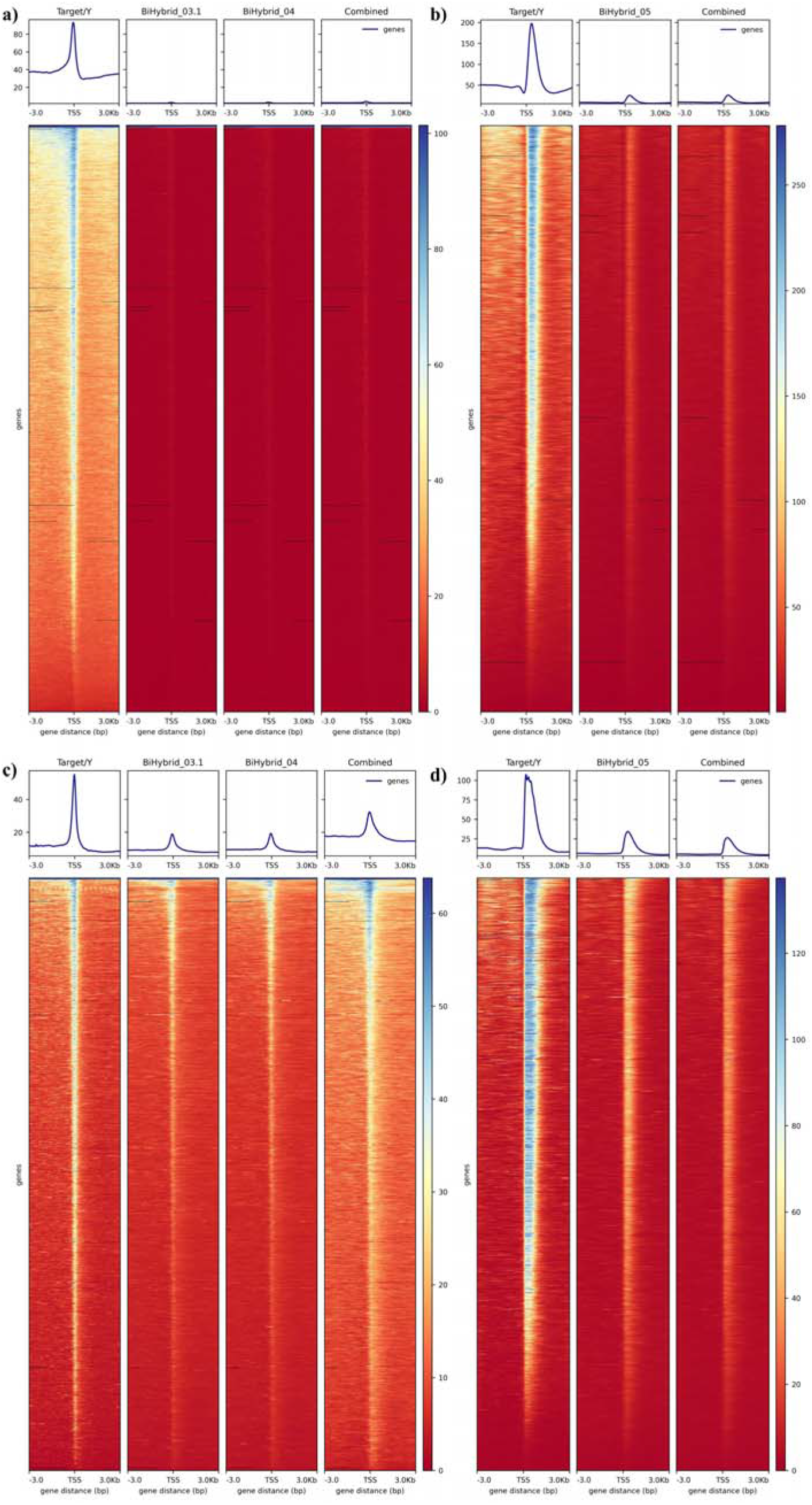
Experimental and predicted ATAC- and ChIP-seq coverage +/- 3 kbp around the TSS. The average experimental coverage (target/y) and predicted coverage, excluding unplaced scaffolds and non-nuclear sequences, in reads per base pair are shown for a) *A. thaliana* ATAC-seq data, b) *A. thaliana* ChIP-seq data, c) *O. sativa* ATAC-seq data and d) *O. sativa* ChIP-seq data. The models used to generate the predictions were the best models using the model architectures BiHybrid_03.1, BiHybrid_04 and the combined model for ATAC-seq and BiHybrid_05 and the combined model for ChIP-seq. The profile plots at the top of each heatmap show the average coverage. The target data was converted from bam to bigwig format using a bin size of 50, so the average coverage of 50 bp subsequences was used. The predictions were converted to bigWig files averaging the individual predictions for the positive and negative strand. See Supplementary Figure S1 for a version of this figure including the predictions for the flagged sequences.

A base-wise F_1_ was calculated to quantify predicted peaks matching experimental peaks (Tab.3). The parameters for peak calling were chosen, so that the peaks called from the mean experimental read coverage of all samples of a given NGS dataset and test species were close to the peaks called for the individual samples (Supplementary Section S1.7). The highest F_1_ score for the ATAC-seq peaks of *A. thaliana* was the combined model’s score of 0.1206. For the ATAC-seq peaks of *O. sativa* the BiHybrid_04 model’s predictions resulted in the highest F_1_ score of 0.3008. In the case of *A. thaliana*, precision, the rate of false negatives, was notably higher than recall, i.e., the true positive rate. This applied to all three tested models. The rate of false negatives was also higher than the true positive rate for the ChIP-seq predictions for *A. thaliana*, but the difference was smaller. For the ATAC-seq predictions of *O. sativa* recall was higher than precision. Precision and recall were balanced for the ChIP-seq predictions of *O. sativa*. The predicted ChIP-seq peaks showed higher F_1_ scores for both test species than the predicted ATAC-seq peaks. The combined model’s F_1_ scores were slightly higher than the BiHybrid_05 model’s scores.

To get a more detailed insight into the model predictions, six zoomed-in example predictions, three per test species, were examined (Fig. 4). The examples were manually selected to present examples for regions with varying levels of prediction quality. By this, we aimed at gaining a deeper understanding of the predictions beyond the quality control using global statistical metrics.

**Figure 4:**
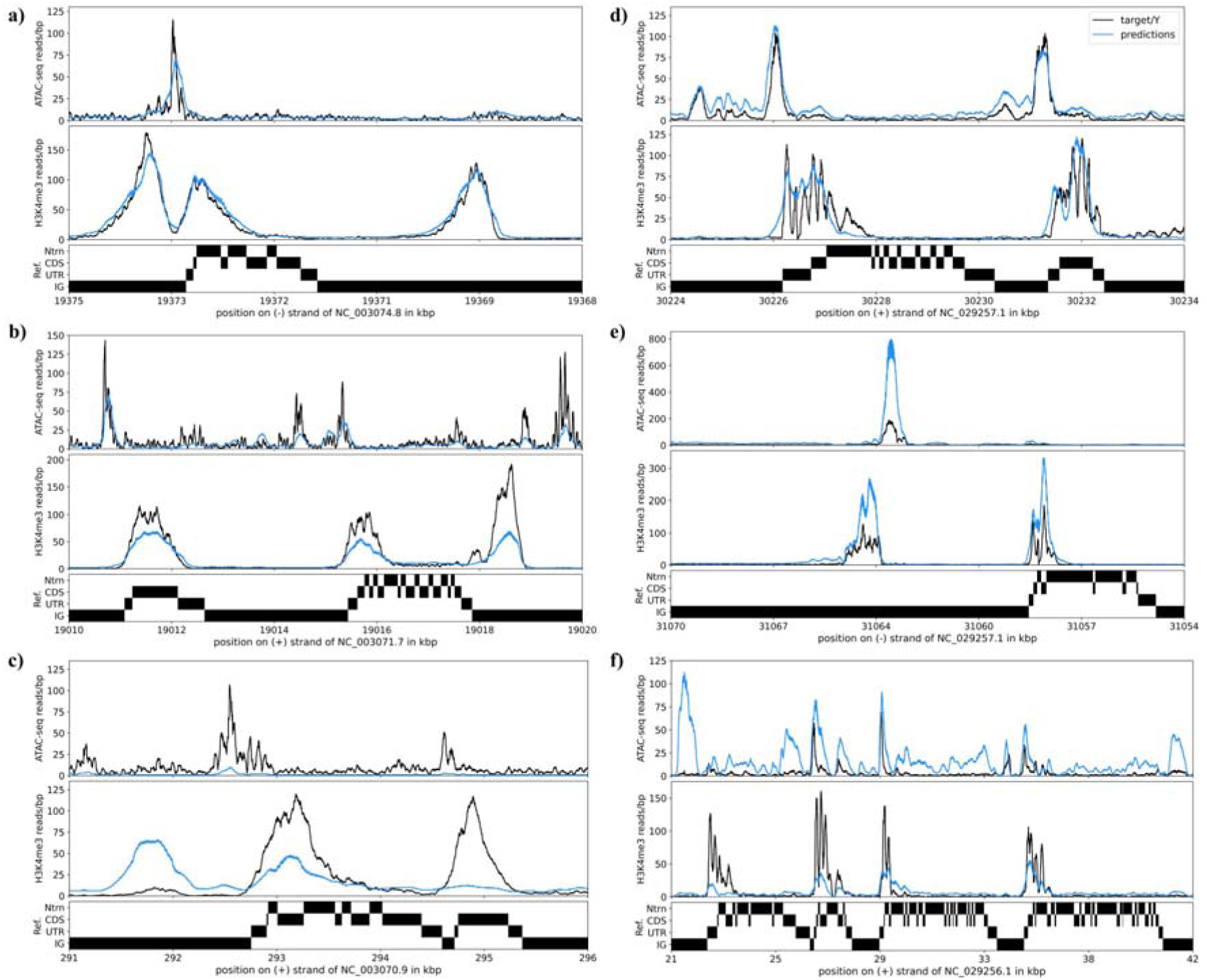
**Example predictions of Predmoter**. Six example regions comparing Predmoter predictions (in blue) to experimental data (target/Y; in black) for the test species a-c) *A. thaliana* and d-f) *O. sativa* in 5’ to 3’ direction are depicted. Three different prediction qualities are shown: a) & d) the predictions and the target data are nearly identical in shape and height, b) & e) the predictions match the target peaks positions but not their exact shape or height, and c) & f) the predicted peaks are in different locations than the target peaks. The combined model was used for the presented predictions. The first section of the plots shows the ATAC-seq reads per bp, the second the ChIP-seq reads per bp. The third section is the reference annotation. It is divided into four categories: intron (Ntrn), coding sequence (CDS), untranslated region (UTR), and intergenic region (IG). The individual predictions for the positive and negative strand are shown respectively. Detailed information on the reference/genome assembly can be found in Supplementary Table S2.

The ChIP-seq predictions for *A. thaliana* had a Pearson’s correlation of 0.98 with no visible outliers compared to the experimental data (Fig. 4a). The ATAC-seq predictions had a Pearson’s correlation of 0.81. The high-quality predictions for *O. sativa* reached a Pearson’s correlation of 0.93 for the ATAC-seq data and 0.85 for the ChIP-seq data (Fig. 4d). The experimental and predicted peaks showed a common pattern of the ATAC-seq peaks around the TSS overlapping the 5’ UTR. They were usually flanked by a H3K4me3 peak downstream of the TSS. Occasionally the ATAC-seq peak was observed between two H3K4me3 peaks, one downstream of the TSS and one upstream of the ATAC-seq peak (Fig. 4a). In the two examples of middling quality the predicted peaks were in the same position but did not show the same shape or amplitude of the experimental data (Fig. 4b & 4e). The predictions for *A. thaliana* in this case were a little lower than the experimental data, reaching a Pearson correlation coefficient of 0.67 for the ATAC-seq data (Fig. 4b). Meanwhile, the predictions for *O. sativa* were notably higher for both ATAC- and ChIP-seq, but the ATAC-seq peak and the two ChIP-seq peaks were in the correct position (Fig. 4e). In instances where Predmoter’s predictions didn’t or only partially matched the experimental data, common occurrences were either the experimental data had a peak in a region, but the predictions did not and vice versa (Fig. 4c & 4f). The ATAC-seq predictions for *A. thaliana* showed no peaks while the experimental data did (Fig. 4c). The H3K4me3 experimental data showed two peaks, one of which Predmoter predicted, the other Predmoter did not predict, resulting in a Pearson’s correlation of 0.28. Additionally, we found one region where a relatively high peak barely supported by the experimental data was predicted (Fig. 4c). The same was observed for the ATAC-seq predictions of *O. sativa* (Fig. 4f). More ATAC-seq peaks than the experimental data would support were predicted. These ATAC-seq predictions also appeared to have more background noise. The ATAC-seq predictions had a Pearson’s correlation of 0.46 (Fig. 4f). On the other hand, the ChIP-seq predictions reached a correlation of 0.75, as the four predicted peaks were in the same location as the experimental peaks but did not show the same amplitude.

Predmoter showed a positive linear correlation between inference times and genome length (Supplementary Section S1.6). Inference took longer the more NGS datasets were predicted simultaneously. Predmoter took 2.84 minutes to predict ATAC- and ChIP-seq data together for *A. thaliana*. For *O. sativa* inference took 11.21 minutes.

## 4. Discussion

The identification of CREs is crucial in any attempts to reconstruct gene regulatory networks. In complex genomes, knowledge is mostly concentrated on coding sequences. Studies focusing on the complex genetic mechanisms behind gene regulation fall behind. The high costs and time investments needed to create ATAC- or ChIP-seq libraries are barriers in the way to unravel the natural variation of gene regulation, especially in non-model plants. We developed Predmoter, a low-threshold, fast and precise DNN that uses the target DNA sequence as input and outputs predicted ATAC-seq and ChIP-seq coverage in human-readable format.

Predmoter used both the positive and negative strand as the model’s input. The ATAC- and ChIP-seq read coverage information was also added to both strands (see 2.2). The advantages were that open chromatin and closed chromatin regions always apply to both strands, so the addition to both strands allowed for built-in data augmentation. The model benefited from the BiLSTM layers extra information (Fig. 2), as they allowed the network to anticipate a gene region when predicting a promoter (Schuster and Paliwal 1997). Also, the bidirectional interpretation of the data was an appropriate inductive bias, given that Predmoter used unstranded data. Even though batch normalization eliminates the need for dropout layers in some cases (Ioffe and Szegedy 2015), adding one dropout layer with a dropout probability of 30 % to Predmoter boosted the predictions (Fig. 2).

The predictions were improved for the ChIP-seq data when predicting both datasets together (Fig. 2). The subsequent slight drop-off in the ATAC-seq predictions could be a result of the network having 26.7 % more ChIP-seq data than ATAC-seq data available for training on after filtering flagged sequences. The network was skewing just lightly to the larger dataset, at least when looking solely at the Pearson correlation coefficients (Fig. 2). The F_1_ scores showed that the predicted ChIP-seq peaks were close to the experimental peaks (Tab. 3). However, the predicted ATAC-seq peaks were not. For the dicot *A. thaliana*, there were many false negative peaks predicted, leading to F_1_ scores from 0.1535 to the highest score of 0.1686 (Tab. 3). This indicates that Predmoter underestimated the number of ATAC-seq peaks or rather the number of ACRs. In the case of the monocot *O. sativa*, Predmoter overestimated the number of ACRs, predicting a high number of false positive peaks. The BiHybrid_04 model had the lowest proportion of false positives leading to the best F_1_ score of 0.3857 for *O. sativa*. H3K4me3 peaks mostly appear in 1000 to 2000 kbp around the TSS including highly conserved gene regions (Santos-Rosa et al. 2002, Benayoun et al. 2014). Even though CREs were shown to be highly conserved within and among plant species, also between monocots and dicots (Yamamoto et al. 2007, Lu et al. 2019), they exhibit heterogeneity. The TATA-box for example, a core promoter element characterized by repeating T and A base pairs (Lifton et al. 1978), was found to be present in 16-22 % of core promoters in eight plant species, in 18 % of the *A. thaliana* and *O. sativa* core promoters (Kumari and Ware 2013). Therefore, the H3K4me3 peaks were probably easier to learn for the network. This could be supported by the models training only on ChIP-seq data or both datasets also reached their highest validation Pearson correlation coefficient faster (see Supplementary Table S6).

**Table 3:** Peak F_1_ statistics. The F_1_ of the predicted peaks versus the experimental peaks was calculated per model, test species and NGS dataset. The predicted and experimental peaks of both strands were compared. Flagged sequences were excluded from peak calling and F_1_ calculation. The number of called peaks is listed in Supplementary Table S9 and the average and median peak lengths in Supplementary Table S10.

| Species<br>(scientific) | ATAC-seq |  |  | ChIP-seq |  |
| --- | --- | --- | --- | --- | --- |
|  | BiHybrid_03.1 | BiHybrid_04 | Combined | BiHybrid_05 | Combined |
| *A. thaliana* | 0.0748 | 0.0786 | 0.1206 | 0.6759 | 0.7033 |
| *O. sativa* | 0.2913 | 0.3008 | 0.1635 | 0.6920 | 0.7049 |

The ATAC-seq coverage enrichment +/- 3 kbp around the TSS of *A. thaliana* and *O. sativa* showed that the predictions of the combined model had a higher amplitude than the BiHybrid_04 model predictions but overestimated the background coverage (Fig. 3). The peak amplitudes depend on the average sequencing depth. Since no adjustments towards sequencing depths were made, the amplitudes were likely technical artifacts. As supported by the F_1_ score of 0.1635 for the combined model predictions for *O. sativa* in comparison to the F_1_ scores of 0.2913 and 0.3008 for the predictions of the other two models (Tab. 3), a higher amplitude did not correlate with more accurate predictions.

In total, the “negative” training set, meaning regions without associated ATAC- or ChIP-seq peaks, was larger than the “positive” one, which could be one reason for some experimentally verified peaks not being predicted (Tab. 3 & Fig. 4c). Predicted peaks in regions missing experimentally verified peaks might appear, because the observed most common pattern was an ATAC-seq peak upstream of a H3K4me3 peak (Fig. 4). The network was likely trying to adhere to that pattern even if the target data did not support it.

Another reason for not predicting experimentally verified peaks or predicting peaks in regions where there are none in the experimental data could be the incompleteness of the experimental data. The experimental data originated from different tissues and was treated differently as well (see Supplementary Table S3). The *A. thaliana* ATAC-seq data for example used DNA extracted from leaves (Lu et al. 2019) and roots (Maher et al. 2018), while the ChIP-seq data used DNA from whole seedlings under cold treatment and after recovery (Xi et al. 2020) and an unknown tissue/treatment. Not all genes are always active in every tissue. The choice of tissues and environmental influences can influence the chromatin makeup of the plants’ DNA. Hence, the experimental data shown was not the ground truth. With the currently publicly available, high-quality data for ATAC-seq and H3K4me3, the possibility of using as many tissues or treatments as possible to train on or even create dedicated models to specific plant tissues like roots is not yet feasible. Some of the available published data was of poor quality. Either, the data did not show the expected pattern around the TSS, or in the case of ATAC-seq data some scaffolded genome assemblies were missing large subsequences of genome information in front of the TSS complicating ATAC-seq data quality control (see 2.1).

Since ATAC-seq and H3K4me3 peaks were seen in this study to be close to each other but only partially overlap, other NGS data showing a more similar pattern to ATAC-seq data could improve the predictions for the ACRs. The nearest options would be DNase-seq (Crawford et al. 2006) or FAIRE- seq (Giresi et al. 2007). Both are less sensitive than ATAC-seq. Another option could be MNase-defined cistrome-Occupancy Analysis (MOA-seq), a high-resolution, high-throughput, and genome-wide strategy to globally identify putative TF-binding sites within ACRs (Savadel et al. 2021). The only hindrance would again be the publicly available high-quality data. For example, MOA-seq is too recent to have large amounts of existing published data. Additional ChIP-seq data, like H3K4me1, H3K27ac or H3K27me3, marking enhancers (Heintzman et al. 2009, Creyghton et al. 2010, Rada- Iglesias et al. 2010) or H3K4me2 marking inactive genes (Santos-Rosa et al. 2002) could be utilized as well. The dicot *A. thaliana* had a different source of error in the form of many non-predicted peaks than the monocot *O. sativa* with many false positive peaks (Tab. 3). Weighting the monocot and dicot data as well as the ATAC- and ChIP-seq data to combat overfitting towards a domain or NGS dataset could improve the predictions. Also, a method of normalizing the NGS read coverage without relying on experimental input data could help the network to focus more on peak positions instead of peak amplitudes. Finally, by incorporating peak caller results like from MACS (Zhang et al. 2008) into the predictive process of Predmoter, the option of a binary classification model could be added. DL was already used with ATAC-seq data and MACS2 to predict regulatory factor binding activity (Hiranuma, Lundberg and Lee 2017), to predict enhancers (Thibodeau et al. 2018) or to optimize ATAC-seq peak calling (Hentges et al. 2022). A drawback to using a peak caller would be the introduction of another abstraction level by using the output of another tool/algorithm. In general, more ATAC-seq data from a wider variety of species and tissues would likely improve Predmoter’s predictions more than additional NGS data.

We are aware that Predmoter is strongly limited by the quality and abundance of ATAC- and ChIP-seq data. However, our framework allows for easy retraining with additional high-quality NGS data. This also includes re-training with selected datasets for tissue- or condition specific treatments. In conclusion, Predmoter will help identifying CREs and so gaining further insight into gene regulatory networks in plants.

## Supplementary material

**Supplementary Figure S1:** Experimental and predicted ATAC- and ChIP-seq coverage +/- 3 kbp around the TSS including flagged sequences

**Supplementary Figure S2:** Benchmarking Helixer’s conversion from fasta to h5 file

**Supplementary Figure S3:** Benchmarking Predmoter’s prediction time of one NGS dataset and conversion time of the output prediction h5 file to bigWig and bedGraph files

**Supplementary Figure S4:** Benchmarking Predmoter’s prediction time of two NGS datasets and conversion time of the output prediction h5 file to bigWig and bedGraph files

**Supplemental Table S1:** One-hot vector encoding used for the input DNA sequence

**Supplemental Table S2:** ATAC-seq or ChIP-seq experimental data origin and plant genome accessions

**Supplemental Table S3:** ATAC-seq or ChIP-seq experimental data tissues and treatments

**Supplemental Table S4:** Model training and validation sets

**Supplemental Table S5:** Model training/Predmoter parameters

**Supplemental Table S6:** Best models per model setup out of the replicate training runs

**Supplemental Table S7:** Pearson’s correlation for ATAC-seq predictions per species and model

**Supplemental Table S8:** Pearson’s correlation for ChIP-seq predictions per species and model

**Supplemental Table S9:** Number of peaks called from the mean experimental read coverage per test species and NGS dataset and of the predicted peaks per model, test species and NGS dataset

**Supplemental Table S10:** Mean and median lengths of peaks called from the mean experimental read coverage per test species and NGS dataset and of the predicted peaks per model, test species and NGS dataset

## Acknowledgements

Computational infrastructure and support were provided by the Centre for Information and Media Technology at Heinrich Heine University Düsseldorf.

## Authors contributions

AD conceived the project idea and supervised the project. AD and FK planned the experiments. FK wrote Predmoter’s code, carried out the experiments, e.g., model training, created the figures and wrote the manuscript. ST and VRD were beta testers and provided expertise on plant promoter and enhancer sequences. ST, US, LAR, APMW and AD edited the manuscript.

## Conflict of interest

The authors declare no conflicts of interest.

## Funding

This work was funded by the Germany’s Excellence Strategy EXC-2048/1 under project ID 390686111 and the Deutsche Forschungsgemeinschaft under project ID 391465903/GRK 2466.

## Notes

https://github.com/weberlab-hhu/Predmoter

### Competing Interest Statement

The authors have declared no competing interest.

## References

1. Andersson, Robin, Gebhard, Claudia, Miguel-Escalada, Irene, Hoof, Ilka, Bornholdt, Jette, Boyd, Mette, et al., ‘An Atlas of Active Enhancers across Human Cell Types and Tissues’, Nature, 507 7493 (2014), 455–61

2. Andrews, S., ‘FastQC A Quality Control Tool for High Throughput Sequence Data’, 2010 <https://www.bioinformatics.babraham.ac.uk/projects/fastqc/> [accessed 23 May 2022]

3. Avsec, Žiga, Agarwal, Vikram, Visentin, Daniel, Ledsam, Joseph R., Grabska-Barwinska, Agnieszka, Taylor, Kyle R., et al., ‘Effective Gene Expression Prediction from Sequence by Integrating Long-Range Interactions’, Nature Methods, 18/10 (2021), 1196–1203

4. Banerji, Julian, Rusconi, Sandro, and Schaffner, Walter, ‘Expression of a β-Globin Gene Is Enhanced by Remote SV40 DNA Sequences’, Cell, 27 (1981), 299–308

5. Benayoun, Bérénice A., Pollina, Elizabeth A., Ucar, Duygu, Mahmoudi, Salah, Karra, Kalpana, Wong, Edith D., et al., ‘H3K4me3 Breadth Is Linked to Cell Identity and Transcriptional Consistency’, Cell, 158/3 (2014), 673–88

6. Bolger, Anthony M., Lohse, Marc, and Usadel, Bjoern, ‘Trimmomatic: A Flexible Trimmer for Illumina Sequence Data’, Bioinformatics, 30/15 (2014), 2114

7. Broad Institute, ‘Picard Toolkit’, *Broad Institute*, GitHub Repository, 2019 <https://broadinstitute.github.io/picard/> [accessed 23 May 2022]

8. Buenrostro, Jason D., Giresi, Paul G., Zaba, Lisa C., Chang, Howard Y., and Greenleaf, William J., ‘Transposition of Native Chromatin for Fast and Sensitive Epigenomic Profiling of Open Chromatin, DNA-Binding Proteins and Nucleosome Position’, Nature Methods, 10/12 (2013), 1213–18

9. Cockerill, Peter N., ‘Structure and Function of Active Chromatin and DNase I Hypersensitive Sites’, The FEBS Journal, 278/13 (2011), 2182–2210

10. Crawford, Gregory E., Holt, Ingeborg E., Whittle, James, Webb, Bryn D., Tai, Denise, Davis, Sean, et al., ‘Genome-Wide Mapping of DNase Hypersensitive Sites Using Massively Parallel Signature Sequencing (MPSS)’, Genome Research, 16/1 (2006), 123

11. Creyghton, Menno P., Cheng, Albert W., Welstead, G. Grant, Kooistra, Tristan, Carey, Bryce W., Steine, Eveline J., et al., ‘Histone H3K27ac Separates Active from Poised Enhancers and Predicts Developmental State’, Proceedings of the National Academy of Sciences of the United States of America, 107/50 (2010), 21931–36

12. Danecek, Petr, Bonfield, James K., Liddle, Jennifer, Marshall, John, Ohan, Valeriu, Pollard, Martin O., et al., ‘Twelve Years of SAMtools and BCFtools’, GigaScience, 10/2 (2021)

13. Dao, Lan T.M., Galindo-Albarrán, Ariel O., Castro-Mondragon, Jaime A., Andrieu-Soler, Charlotte, Medina-Rivera, Alejandra, Souaid, Charbel, et al., ‘Genome-Wide Characterization of Mammalian Promoters with Distal Enhancer Functions’, Nature Genetics, 49/7 (2017), 1073–81

14. Diao, Yarui, Fang, Rongxin, Li, Bin, Meng, Zhipeng, Yu, Juntao, Qiu, Yunjiang, et al., ‘A Tiling-Deletion-Based Genetic Screen for Cis-Regulatory Element Identification in Mammalian Cells’, Nature Methods, 14/6 (2017), 629–35

15. Dynan, William S., and Tjian, Robert, ‘Control of Eukaryotic Messenger RNA Synthesis by Sequence-Specific DNA-Binding Proteins’, Nature, 316/6031 (1985), 774–78

16. Engreitz, Jesse M., Haines, Jenna E., Perez, Elizabeth M., Munson, Glen, Chen, Jenny, Kane, Michael, et al., ‘Local Regulation of Gene Expression by LncRNA Promoters, Transcription and Splicing’, Nature, 539/7629 (2016), 452–55

17. Epstein, Wolfgang, and Beckwith, Jonathan R, ‘Regulation of Gene Expression’, Annual Review of Biochemistry, 37/1 (1968), 411–36

18. Ewels, Philip, Magnusson, Måns, Lundin, Sverker, and Käller, Max, ‘MultiQC: Summarize Analysis Results for Multiple Tools and Samples in a Single Report’, Bioinformatics, 32/19 (2016), 3047– 48

19. Falcon, William, ‘Pytorch Lightning: GitHub’, 2019 <https://github.com/PyTorchLightning> [accessed 19 April 2022]

20. Giresi, Paul G., Kim, Jonghwan, McDaniell, Ryan M., Iyer, Vishwanath R., and Lieb, Jason D., ‘FAIRE (Formaldehyde-Assisted Isolation of Regulatory Elements) Isolates Active Regulatory Elements from Human Chromatin’, Genome Research, 17/6 (2007), 877–85

21. Glorot, Xavier, Bordes, Antoine, and Bengio, Yoshua, ‘Deep Sparse Rectifier Neural Networks’ (2011), 315–23

22. Gross, David S, and Garrard, William T, ‘Nuclease Hypersensitive Sites in Chromatin’, Annual Review of Biochemistry, 57, 1988, 159–97

23. Heintzman, Nathaniel D., Hon, Gary C., Hawkins, R. David, Kheradpour, Pouya, Stark, Alexander, Harp, Lindsey F., et al., ‘Histone Modifications at Human Enhancers Reflect Global Cell-Type-Specific Gene Expression’, Nature, 459/7243 (2009), 108–12

24. Hentges, Lance D., Sergeant, Martin J., Cole, Christopher B., Downes, Damien J., Hughes, Jim R., and Taylor, Stephen, ‘LanceOtron: A Deep Learning Peak Caller for Genome Sequencing Experiments’, Bioinformatics, 38/18 (2022), 4255–63

25. Hiranuma, Naozumi, Lundberg, Scott, and Lee, Su-In, ‘DeepATAC: A Deep-Learning Method to Predict Regulatory Factor Binding Activity from ATAC-Seq Signals’, BioRxiv, 2017

26. Hochreiter, Sepp, and Schmidhuber, Jürgen, ‘Long Short-Term Memory’, Neural Computation, 9/8 (1997), 1735–80

27. Holst, Felix, Bolger, Anthony, Günther, Christopher, Maß, Janina, Triesch, Sebastian, Kindel, Felicitas, et al., ‘Helixer–de Novo Prediction of Primary Eukaryotic Gene Models Combining Deep Learning and a Hidden Markov Model’, BioRxiv, 2023

28. Ioffe, Sergey, and Szegedy, Christian, ‘Batch Normalization: Accelerating Deep Network Training by Reducing Internal Covariate Shift’, in Proceedings of the 32nd International Conference on Machine Learning (2015), 448–56

29. Ippen, Karin, Miller, Jeffrey H., Scaife, John, and Beckwith, Jonathan, ‘New Controlling Element in the Lac Operon of E. Coli’, Nature, 217/5131 (1968), 825–27

30. Jacob, F, Ullman, A, and Monod, J, ‘Le Promoteur, Élément Génétique Nécessaire à l’expression d’un Opéron’, CR Acad. Sci.(Paris), 258 (1964), 3125–28

31. Johnson, David S., Mortazavi, Ali, Myers, Richard M., and Wold, Barbara, ‘Genome-Wide Mapping of in Vivo Protein-DNA Interactions’, Science, 316/5830 (2007), 1497–1502

32. Kim, Jung-whan, Zeller, Karen I., Wang, Yunyue, Jegga, Anil G., Aronow, Bruce J., O’Donnell, Kathryn A., et al., ‘Evaluation of Myc E-Box Phylogenetic Footprints in Glycolytic Genes by Chromatin Immunoprecipitation Assays’, Molecular and Cellular Biology, 24/13 (2004), 5923– 36

33. Kim, Tae Kyung, Hemberg, Martin, Gray, Jesse M., Costa, Allen M., Bear, Daniel M., Wu, Jing, et al., ‘Widespread Transcription at Neuronal Activity-Regulated Enhancers’, Nature, 465/7295 (2010), 182–87

34. Kingma, Diederik P., and Ba, Jimmy Lei, ‘Adam: A Method for Stochastic Optimization’, 3rd International Conference on Learning Representations, ICLR 2015 - Conference Track Proceedings, 2014

35. Kumari, Sunita, and Ware, Doreen, ‘Genome-Wide Computational Prediction and Analysis of Core Promoter Elements across Plant Monocots and Dicots’, PLOS ONE, 8/10 (2013), e79011

36. LeCun, Yann, and Bengio, Yoshua, ‘Convolutional Networks for Images, Speech, and Time-Series’, The Handbook of Brain Theory and Neural Networks, 1995, 255–58

37. LeCun, Yann, Boser, Bernhard, Denker, John, Henderson, Donnie, Howard, R, Hubbard, Wayne, et al., ‘Handwritten Digit Recognition with a Back-Propagation Network’, Advances in Neural Information Processing Systems, 2 (1989)

38. Lifton, R. P., Goldberg, M. L., Karp, R. W., and Hogness, D. S., ‘The Organization of the Histone Genes in Drosophila Melanogaster: Functional and Evolutionary Implications’, Cold Spring Harbor Symposia on Quantitative Biology, 42/2 (1978), 1047–51

39. Lu, Zefu, Marand, Alexandre P., Ricci, William A., Ethridge, Christina L., Zhang, Xiaoyu, and Schmitz, Robert J., ‘The Prevalence, Evolution and Chromatin Signatures of Plant Regulatory Elements’, Nature Plants, 5/12 (2019), 1250–59

40. Maher, Kelsey A., Bajic, Marko, Kajala, Kaisa, Reynoso, Mauricio, Pauluzzi, Germain, West, Donnelly A., et al., ‘Profiling of Accessible Chromatin Regions across Multiple Plant Species and Cell Types Reveals Common Gene Regulatory Principles and New Control Modules’, The Plant Cell, 30/1 (2018), 15–36

41. Md, Vasimuddin, Misra, Sanchit, Li, Heng, and Aluru, Srinivas, ‘Efficient Architecture-Aware Acceleration of BWA-MEM for Multicore Systems’, Proceedings - 2019 IEEE 33rd International Parallel and Distributed Processing Symposium, IPDPS 2019, 2019, 314–24

42. Oubounyt, Mhaned, Louadi, Zakaria, Tayara, Hilal, and To Chong, Kil, ‘Deepromoter: Robust Promoter Predictor Using Deep Learning’, Frontiers in Genetics, 10/APR (2019), 286

43. Paszke, Adam, Gross, Sam, Massa, Francisco, Lerer, Adam, Bradbury, James, Chanan, Gregory, et al., ‘PyTorch: An Imperative Style, High-Performance Deep Learning Library’, Advances in Neural Information Processing Systems, 32 (2019)

44. Rada-Iglesias, Alvaro, Bajpai, Ruchi, Swigut, Tomek, Brugmann, Samantha A., Flynn, Ryan A., and Wysocka, Joanna, ‘A Unique Chromatin Signature Uncovers Early Developmental Enhancers in Humans’, Nature, 470/7333 (2010), 279–83

45. Ramírez, Fidel, Ryan, Devon P., Grüning, Björn, Bhardwaj, Vivek, Kilpert, Fabian, Richter, Andreas S., et al., ‘DeepTools2: A next Generation Web Server for Deep-Sequencing Data Analysis’, Nucleic Acids Research, 44/W1 (2016), W160–65

46. Robertson, Gordon, Hirst, Martin, Bainbridge, Matthew, Bilenky, Misha, Zhao, Yongjun, Zeng, Thomas, et al., ‘Genome-Wide Profiles of STAT1 DNA Association Using Chromatin Immunoprecipitation and Massively Parallel Sequencing’, Nature Methods, 4/8 (2007), 651–57

47. Russell, Stuart J., and Norvig, Peter, Artificial Intelligence: A Modern Appoach., Global Edition. (2016)

48. de Santa, Francesca, Barozzi, Iros, Mietton, Flore, Ghisletti, Serena, Polletti, Sara, Tusi, Betsabeh Khoramian, et al., ‘A Large Fraction of Extragenic RNA Pol II Transcription Sites Overlap Enhancers’, PLOS Biology, 8/5 (2010), e1000384

49. Santos-Rosa, Helena, Schneider, Robert, Bannister, Andrew J., Sherriff, Julia, Bernstein, Bradley E., Emre, N. C.Tolga, et al., ‘Active Genes Are Tri-Methylated at K4 of Histone H3’, Nature, 419/6905 (2002), 407–11

50. Savadel, Savannah D., Hartwig, Thomas, Turpin, Zachary M., Vera, Daniel L., Lung, Pei Yau, Sui, Xin, et al., ‘The Native Cistrome and Sequence Motif Families of the Maize Ear’, PLOS Genetics, 17/8 (2021), e1009689

51. Schulz, Hannes, and Behnke, Sven, ‘Deep Learning: Layer-Wise Learning of Feature Hierarchies’, *KI- Kunstliche Intelligenz*, 26/4 (2012), 357–63

52. Schuster, Mike, and Paliwal, Kuldip K., ‘Bidirectional Recurrent Neural Networks’, IEEE Transactions on Signal Processing, 45/11 (1997), 2673–81

53. Shahmuradov, Ilham A., Umarov, Ramzan K., and Solovyev, Victor V., ‘TSSPlant: A New Tool for Prediction of Plant Pol II Promoters’, Nucleic Acids Research, 45/8 (2017), e65

54. Shujaat, Muhammad, Lee, Seung Beop, Tayara, Hilal, and Chong, Kil To, ‘Cr-Prom: A Convolutional Neural Network-Based Model for the Prediction of Rice Promoters’, IEEE Access, 9 (2021), 81485–91

55. Song, Lingyun, Zhang, Zhancheng, Grasfeder, Linda L., Boyle, Alan P., Giresi, Paul G., Lee, Bum Kyu, et al., ‘Open Chromatin Defined by DNaseI and FAIRE Identifies Regulatory Elements That Shape Cell-Type Identity’, Genome Research, 21/10 (2011), 1757–67

56. Stiehler, Felix, Steinborn, Marvin, Scholz, Stephan, Dey, Daniela, Weber, Andreas P.M., and Denton, Alisandra K., ‘Helixer: Cross-Species Gene Annotation of Large Eukaryotic Genomes Using Deep Learning’, Bioinformatics, 36/22–23 (2021), 5291–98

57. Struhl, Kevin, ‘Yeast Transcriptional Regulatory Mechanisms’, Annual Review of Genetics, 29 (1995), 651–74

58. Thibodeau, Asa, Uyar, Asli, Khetan, Shubham, Stitzel, Michael L., and Ucar, Duygu, ‘A Neural Network Based Model Effectively Predicts Enhancers from Clinical ATAC-Seq Samples’, Scientific Reports, 8/1 (2018), 1–15

59. Umarov, Ramzan Kh, and Solovyev, Victor V., ‘Recognition of Prokaryotic and Eukaryotic Promoters Using Convolutional Deep Learning Neural Networks’, PLOS ONE, 12/2 (2017), e0171410

60. Umarov, Ramzan, Kuwahara, Hiroyuki, Li, Yu, Gao, Xin, and Solovyev, Victor, ‘Promoter Analysis and Prediction in the Human Genome Using Sequence-Based Deep Learning Models’, Bioinformatics, 35/16 (2019), 2730–37

61. Wang, Ying, Peng, Qinke, Mou, Xu, Wang, Xinyuan, Li, Haozhou, Han, Tian, et al., ‘A Successful Hybrid Deep Learning Model Aiming at Promoter Identification’, BMC Bioinformatics, 23/1 (2022), 1–20

62. Xi, Yanpeng, Park, Sung Rye, Kim, Dong Hwan, Kim, Eun Deok, and Sung, Sibum, ‘Transcriptome and Epigenome Analyses of Vernalization in Arabidopsis Thaliana’, The Plant Journal, 103/4 (2020), 1490–1502

63. Yamamoto, Yoshiharu Y., Ichida, Hiroyuki, Matsui, Minami, Obokata, Junichi, Sakurai, Tetsuya, Satou, Masakazu, et al., ‘Identification of Plant Promoter Constituents by Analysis of Local Distribution of Short Sequences’, BMC Genomics, 8/1 (2007), 1–23

64. Zhang, Yong, Liu, Tao, Meyer, Clifford A., Eeckhoute, Jérôme, Johnson, David S., Bernstein, Bradley E., et al., ‘Model-Based Analysis of ChIP-Seq (MACS)’, Genome Biology, 9/9 (2008), 1–9

65. Zheng, Linlin, McMullen, Michael D., Bauer, Eva, Schön, Chris Carolin, Gierl, Alfons, and Frey, Monika, ‘Prolonged Expression of the BX1 Signature Enzyme Is Associated with a Recombination Hotspot in the Benzoxazinoid Gene Cluster in Zea Mays’, Journal of Experimental Botany, 66/13 (2015), 3917–30

