## Supplementary material for "Predmoter - Cross-species prediction of plant promoter and enhancer regions"

### Supplemental material for

### Predmoter - Cross-species prediction of plant promoter and enhancer associated NGS data

---

### S1 Supplemental Methods

#### S1.1 Data

The plant genomes were acquired from NCBI's RefSeq or GenBank. The ATAC- and ChIP-seq data is publicly available data acquired from the NCBI's SRA (Tab. S2). The different plant tissues and treatments used in the ATAC- and ChIP-seq experiments are listed if available (Tab. S3).

#### S1.2 Architecture

Predmoter uses custom padding formulas to ensure sequence length divisibility and tensor shape consistency. The padding formulas are adapted from the PyTorch documentation of the formulas to calculate the output sequence length of a one-dimensional convolution (Equation S1) and a one-dimensional transposed convolution (Equation S3) respectively. The initial input sequence length, default is 21384 bp, needs to be divisible by the chosen stride (referred to as step in Predmoter). For multiple convolutional layers the sequence length needs to be divisible by the chosen stride to the power of the chosen number of convolutional layers. Since the convolutional and transposed convolutional layers don't evenly divide the given sequence length, i.e., a sequence length of 21384 and a stride of 2 should result in an output sequence length of 10692, the padding formulas (Equation S2 & S4) are used to control the amount of padding applied to both sides of the input, default is adding zeros, to ensure even division. The output padding for the transposed convolutional layers is added to one side of the output; the default is also to add zeros.

**Equation S1: Calculation of the output sequence length of a one-dimensional convolutional layer**
The output sequence length ($L_{out}$) is calculated using the input variables: padding, dilation, kernel size, stride, and the input sequence length ($L_{in}$) of the one-dimensional convolutional layer. The brackets indicate to round the result down (https://pytorch.org/docs/stable/generated/torch.nn.Conv1d.html).

$$L_{out} = \left\lfloor \frac{L_{in} + 2 * padding - dilation * (kernelsize - 1) - 1}{stride} + 1 \right\rfloor$$

**Equation S2: Calculation of padding applied to a one-dimensional convolutional layer**
Padding is calculated using the variables: output sequence length ($L_{out}$), dilation, kernel size, stride, and the input sequence length ($L_{in}$) of the one-dimensional convolutional layer, where the desired $L_{out}$ is: $L_{out} = \frac{L_{in}}{stride}$. The brackets indicate to round the result up.

$$padding = \left\lceil \frac{(L_{out} - 1) * stride - L_{in} + dilation * (kernelsize - 1) + 1}{2} \right\rceil$$

**Equation S3: Calculation of the output sequence length of a one-dimensional transposed convolutional layer**
The output sequence length ($L_{out}$) is calculated using the input variables padding, output padding, dilation, kernel size, stride, and the input sequence length ($L_{in}$) of the transposed convolutional layer (https://pytorch.org/docs/stable/generated/torch.nn.ConvTranspose1d.html).

$$L_{out} = (L_{in} - 1) * stride - 2 * padding + dilation * (kernelsize - 1) + output\_padding + 1$$

**Equation S4: Calculation of padding applied to a one-dimensional transposed convolutional layer**
Padding is calculated using the variables: output sequence length ($L_{out}$), output padding, dilation, kernel size, stride, and the input sequence length ($L_{in}$), where the desired $L_{out}$ is: $L_{out} = L_{in} * \text{stride}$. The

combination of an even kernel size and a stride of 1 is currently not possible due to a limitation in PyTorch.

$$padding = \frac{(L_{in} - 1) * stride - L_{out} + dilation * (kernel\_size - 1) + output\_padding + 1}{2}$$

where:

$$output\_padding = \begin{cases} 0, & if \quad 2 \mid (kernel\_size + stride) \\ 1, & otherwise \end{cases}$$

#### S1.3 Training details

The models were trained and tested on a server with an Intel(R) Xeon(R) CPU E5-2640v4 (Broadwell) @ 2.40 GHz and a Nvidia GeForce GTX 1080 Ti GPU (11 Gb memory). The software versions were CUDA 11.7.1, cuDNN 8.7.0 and Python 3.8.3. The relevant Python package versions were PyTorch 2.0.1, Lightning 2.0.4, Helixer version 0.3.2 and Predmoter version 0.3.2 were used. The exact versions of the other packages used can be found in: https://github.com/weberlab-hhu/Predmoter/blob/main/training_package_versions_freeze.txt. The same server and setup were used for generating the predictions for *Arabidopsis thaliana* and *Oryza sativa*.

Three replicates per model setup were trained utilizing three different seeds: 132709648, 961333724 and 4227086911. The species selection for each model setup is listed in Table S4. The exact training parameters can be found in Table S5. The models were trained until convergence, meaning until the listed “stop-quantity” stopped improving for the set number of epochs (patience). The best models, the models with the highest Pearson’s r value for the validation set are listed in Table S6. These models were used to compare the target data to the model’s predictions for each species from the training, validation, and test set. Testing was performed with four workers/CPUs, one GPU and a batch size of 200. The tabular results are listed in Tables S7 and S8. Predictions on the test set were also generated with four workers/CPUs, one GPU and a batch size of 200. Direct conversion from the h5 output file to a or multiple bigwig files was chosen. The predictions for the test species including the flagged regions are shown in Figure S1.

#### S1.4 Log files

Predmoter creates one to two output log files. The standard log file, <prefix>_predmoter.log, contains important information about parameters chosen and time taken for training, testing, or predicting as well as file conversion steps. This is also the output log of convert2coverage.py. The training metrics log file, <prefix>_metrics.log, is only created during training and consists of the epoch numbers, average validation loss, average validation “accuracy”, average training loss, and average training “accuracy” in that order. The test metrics log file, <prefix>_test_metrics.log, is only created during testing and contains either the total average validation loss and average validation “accuracy” per h5 file/plant species as well as the average validation loss and average validation “accuracy” per NGS dataset or, when just one NGS dataset was used to train the model, just the metrics for this dataset. Predmoter never deletes these log files, but instead appends to them, meaning all information about training, resuming training, testing, and predicting stay together in one file.

#### S1.5 Filtering flagged sequences

A naïve filtering approach was used to reduce the noise in the dataset. ATAC-seq data shows very high coverage for non-nuclear sequences. The transposase cuts primarily open chromatin (Buenrostro et al. 2013) and as such also the chloroplast and mitochondrial genomes. When the organelles were not completely removed before the experiment, the data contains noise in the form of notably higher coverage in these regions. Unplaced scaffolds were also observed to contribute to this noise during the data quality control steps. Therefore, unplaced scaffolds and non-nuclear sequences were flagged starting with the model architecture BiHybrid_04 (Tab. S5). Assemblies on scaffold or contig level, *Bigelowiella natans*, *Eragrostis nindensis*, *Marchantia polymorpha*, *Oropetium thomaeum*, *Pyrus x bretschneiderii* and *Spirodela polyrhiza,* were not flagged. The flagged sequences were filtered out. The information about the assembly accessions of the unplaced scaffolds and non-nuclear sequences was extracted from the sequence report jsonl files available at the NCBI's RefSeq or GenBank and added to the h5 file (under "data/blacklist") via add_blacklist.py in "side_scripts".

#### S1.6 Benchmarking

Benchmarking was performed on a machine with an Intel(R) Xeon(R) CPU W-2125 @ 4.00 GHz and a Nvidia GeForce GTX 1050 Ti GPU (4 Gb memory). The software versions were CUDA 11.5, cuDNN 8.9.5 and Python 3.10.12. The relevant Python package versions were PyTorch 2.0.1, Lightning 2.0.8, Helixer version 0.3.2 and Predmoter version 0.3.2 were used. The exact versions of the other packages used can be found in: https://github.com/weberlab-hhu/Predmoter/blob/main/benchmarking_package_versions_freeze.txt.

Depending on the model used, there is always a slight fluctuation in the prediction and conversion to bigWig or bedGraph files. Two different models BiHybrid_04 and the combined model were used, as predicting two datasets increases the computing time. Three benchmarking figures are shown. The first shows benchmarking Helixer's conversion from fasta to h5 files (Fig. S2). The second shows benchmarking inference and converting these into bigWig and bedGraph files using BiHybrid_04, a model only trained on and able to predict ATAC-seq data (Fig. S3). The final figure shows benchmarking inference and converting these into bigWig and bedGraph files using the combined model trained on and able to predict ATAC- and ChIP-seq data (Fig. S4). Some genome assemblies were highly fragmented, on contig or scaffold level, increasing the number of subsequences. For example, the genome assembly of *Arabidopsis thaliana* wasn't highly fragmented, the genome size being 119.7 Mbp and the number of 21384 bp subsequences of the h5 file created by Helixer was 11202. The genome of *B. natans* was highly fragmented, the genome size being 91.4 Mbp, but the h5 file contained 13390 subsequences. Since inference and conversion to bigWig or bedGraph files is dependent on the amount of data, so the number of subsequences, that was used to quantify the wall time (Fig. S3 & S4).

#### S1.7 Peak calling

MACS3 (Zhang et al. 2008) was used for calling peaks. Flagged sequences were excluded from the calculations (see Section S1.5). The model trained and was tested on the mean experimental read coverage of all SRA samples, e.g., all 5 ATAC-seq samples of *A. thaliana* (see Table S2). Peaks were called from bedGraph files. The experimental peaks, i.e., the mean experimental read coverage, were called per test species and NGS dataset. The predicted peaks of 4 different models, BiHybrid_031, BiHybrid_04, BiHybrid_05 and the combined model, were called per test species. For ATAC-seq "bdgpeakcall" was used and for ChIP-seq "bdgbroadcall". The minimum peak length was set to the median insert size and the maximum gap size to the mean read length of all samples of one species and one NGS dataset. The median insert size and mean read lengths of each sample ID was collected with Qualimap (Okonechnikov, Conesa and García-Alcalde 2016). For the ATAC-seq data of *A. thaliana* the minimum read length was set to 85 and the maximum gap size to 35, for *O. sativa* the minimum read length was set to 60 and the maximum gap size to 70. For the ChIP-seq data of *A. thaliana* the minimum read length was set to 180 and the maximum gap size to 60, for *O. sativa* the minimum read length was set to 145 and the maximum gap size to 95. The maximum linking gap of "bdgbroadcall" was set to four times the minimum peak length. The cutoff parameter of "bdgbroadcall" had no influence on the number of peaks getting called and was set to 120. To find the optimal cutoff ("bdgpeakcall") or linking cutoff ("bdgbroadcall") an area under the precision-recall curve (AUPRC) was calculated. The optimal thresholds for these two parameters were defined as the smallest difference between precision and recall. Peaks called from the bedGraph files of mean experimental read coverage were compared with peaks called from the individual SRA sample bam files. The peaks of the individual samples were called with MACS3's "callpeak" command, a sophisticated algorithm calling peaks from alignment results. The parameters format BAMPE, a mappable genome size of 119,482,990 bp for *A. thaliana* and 374,305,350 bp for *O. sativa*, a q-value of 0.01 for ATAC-seq data and a q-value of 0.05 for ChIP-seq data and keeping all duplicate tags were utilized. Additionally, the calling broad peaks option was added for calling ChIP-seq peaks. 20 cutoff values were tested, a range of 5 to 100 in intervals of 5. The same values were used for testing the linking cutoff. The cutoff for calling ATAC-seq peaks resulting in the smallest difference between precision and recall was 10 for both test species. The optimal linking cutoff for calling ChIP-seq peaks 20 for *A. thaliana* and 15 for *O. sativa*. These values were also used for calling the predicted peaks. The number of called peaks is listed in Table S9 and the average and median peak lengths in Table S10.

##### S1.8 Figure creation

The taxonomy tree in Figure 2 was created with the NCBI's Taxonomy Common Tree application and visualized using iTOL (Letunic and Bork 2021). The heatmaps were created with Matplotlib (Hunter 2007) and Seaborn (Waskom 2021). The plots in Figure 3 and Figure S1 were created with deepTools (Ramírez et al. 2016). The plots in Figure 4 were created with Matplotlib.

### S2 Supplemental Figures

#### S2.1 Alternative figures

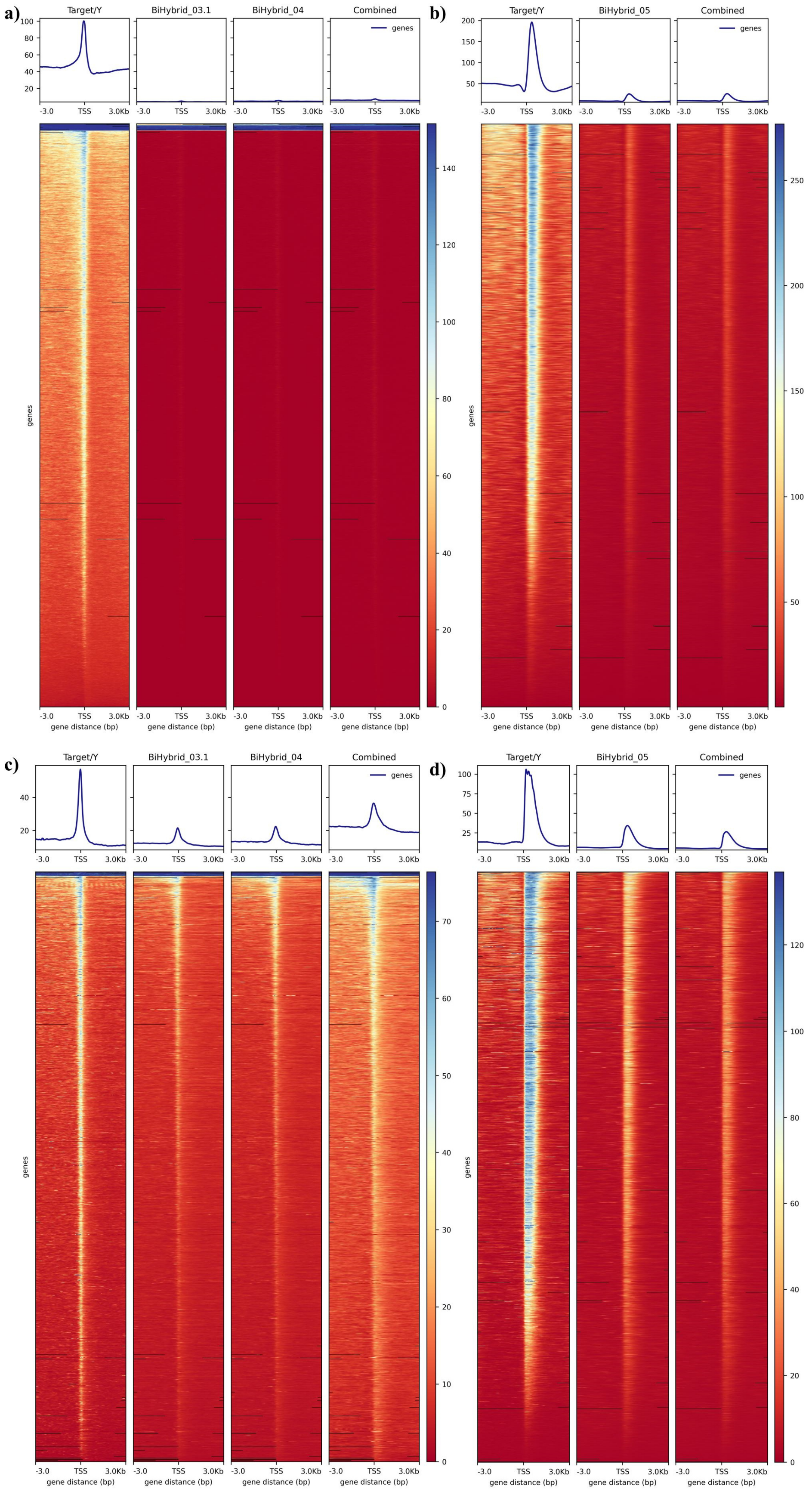

**Figure S1: Experimental and predicted ATAC- and ChIP-seq coverage +/- 3 kbp around the TSS.** The average experimental coverage (target/y) and predicted coverage in reads per base pair are shown for a) *A. thaliana* ATAC-seq data, b) *A. thaliana* ChIP-seq data, c) *O. sativa* ATAC-seq data and d) *O. sativa* ChIP-seq data. The models used to generate the predictions were the best models using the model architectures BiHybrid_03.1, BiHybrid_04 and the combined model for ATAC-seq and BiHybrid_05 and the combined model for ChIP-seq. The profile plots at the top of each heatmap show the average coverage. The target data was converted from bam to bigwig format using a bin size of 50, so the average coverage of 50 bp subsequences was used.

#### S2.2 Benchmarking

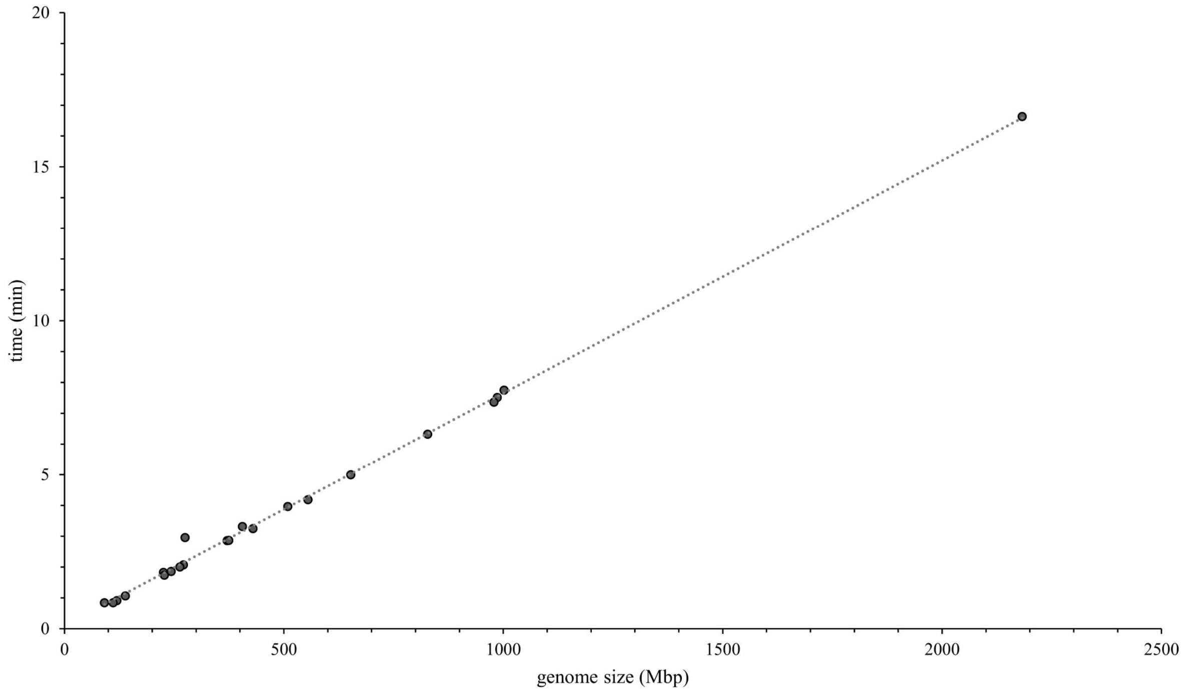

**Figure S2: Benchmarking conversion from fasta to h5 file.** Helixer's wall time for converting fasta to h5 files in minutes for all the species/genome assemblies used in this study. The gapped genome size including unplaced scaffolds in Mbp is shown on the x-axis. The individual data points and a linear trend line are depicted.

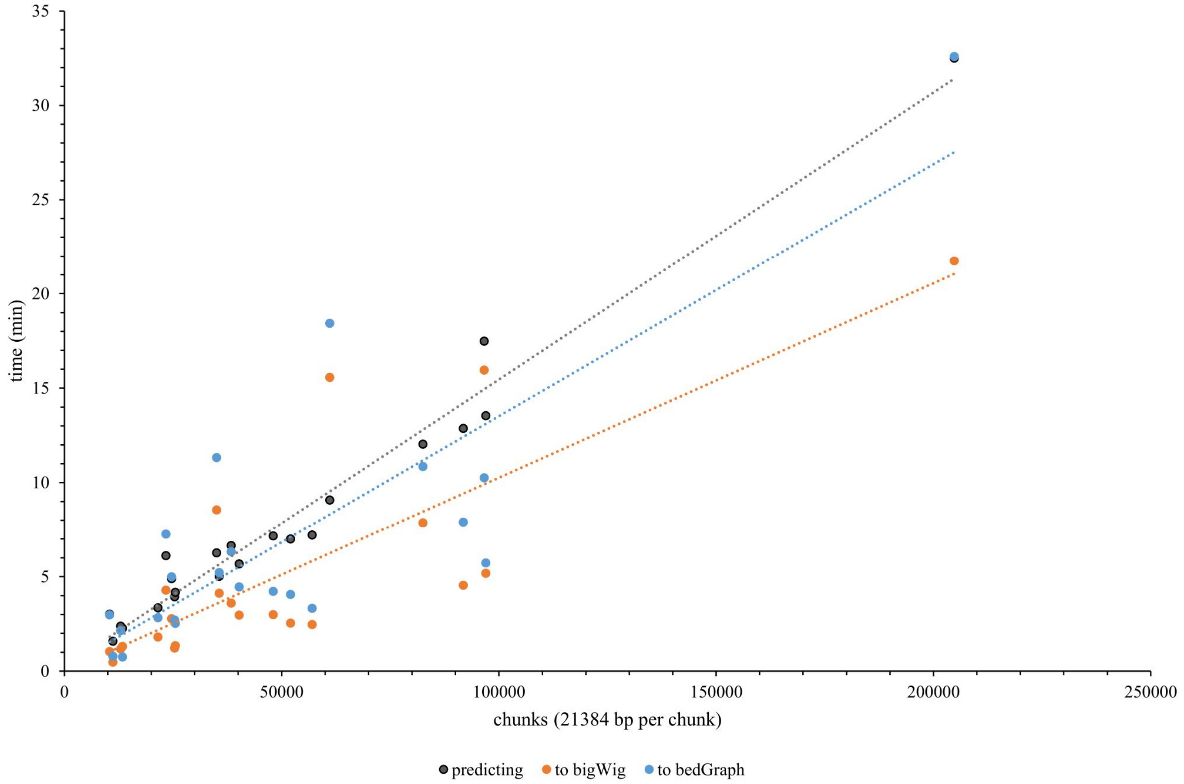

**Figure S3: Benchmarking 1.** The prediction time (black) and prediction h5 file conversion time to bigWig (orange) and bedGraph (blue) files for ATAC-seq read coverage are depicted. The model used for predicting was the BiHybrid_04 model. The wall time in minutes is shown on the y-axis and the chunk number, one chunk is 21384 bp long, on the x-axis. The individual data points and a linear trend line are displayed.

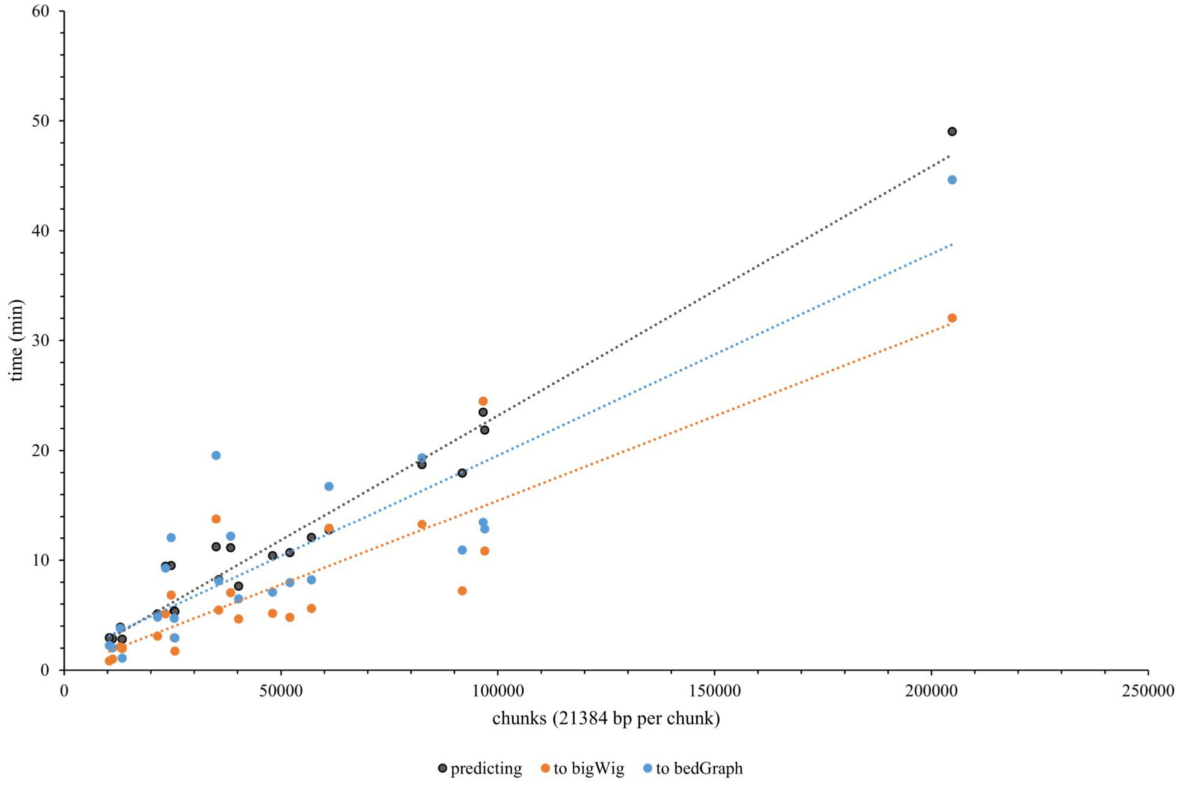

**Figure S4: Benchmarking 2.** The prediction time (black) and prediction h5 file conversion time to bigWig (orange) and bedGraph (blue) files for ATAC- and ChIP-seq read coverage are depicted. The model used for predicting was the combined model. The wall time in minutes is shown on the y-axis and the chunk number, one chunk is 21384 bp long, on the x-axis. The individual data points and a linear trend line are displayed.

### S3 Supplemental Tables

#### S3.1 Data

**Table S1: Nucleotide encoding.** The bases and possible other notations aside from C, A, T or G and the corresponding one-hot vector encoding used for the input DNA sequence are listed.

| Base | Encoding |
|---|---|
| C | [1., 0., 0., 0.] |
| A | [0., 1., 0., 0.] |
| T | [0., 0., 1., 0.] |
| G | [0., 0., 0., 1.] |
| Y | [0.5, 0., 0.5, 0.] |
| R | [0., 0.5, 0., 0.5] |
| W | [0., 0.5, 0.5, 0.] |
| S | [0.5, 0., 0., 0.5] |
| K | [0., 0., 0.5, 0.5] |
| M | [0.5, 0.5, 0., 0.] |
| D | [0., 0.33, 0.33, 0.33] |
| V | [0.33, 0.33, 0., 0.33] |
| H | [0.33, 0.33, 0.33, 0.] |
| B | [0.33, 0., 0.33, 0.33] |
| N | [0.25, 0.25, 0.25, 0.25] |

**Table S2: Data origin.** The table contains the species, the BioProject and the linked SRA accessions of either the ATAC-seq or ChIP-seq experiment passing data preprocessing and quality control, the genome assembly, and the split into training (train), validation (val) or test species.

| Species (scientific) | ATAC-seq (SRA and BioProject accessions) | ChIP-seq (SRA and BioProject accessions) | Genome (RefSeq/GenBank accession) | Split |
|---|---|---|---|---|
| *Arabidopsis thaliana* | PRJNA394532:<br>SRS2357129<br>SRS2357131<br>SRS2357132<br>PRJNA527732:<br>SRS4500485<br>SRS4500486 | PRJNA408288:<br>SRR6057430<br>SRR6057434<br>PRJNA535479:<br>SRS4672527<br>SRS4672528<br>SRS4672529 | GCF_000001735.4 | test |
| *Bigelowiella natans* | PRJNA753294:<br>SRS9735275 | | GCA_000320545.1 | train |
| *Brachypodium distachyon* | PRJNA661629:<br>SRS7327661<br>SRS7327662<br>SRS7327689<br>SRS7327690 | PRJNA661629:<br>SRS7327667<br>SRS7327668<br>SRS7327669<br>SRS7327670 | GCF_000005505.3 | train |
| *Brassica napus* | PRJNA808238:<br>SRS12055968<br>SRS12055970 | PRJNA687926:<br>SRS7933167 | GCF_020379485.1 | train |
| *Brassica oleracea* | | PRJNA687926:<br>SRS7933149 | GCA_900416815.2 | train |
| *Brassica rapa* | | PRJNA687926:<br>SRS7933147 | GCA_016163755.1 | train |
| *Chlamydomonas reinhardtii* | | PRJNA681680:<br>SRR13170450 | GCF_000002595.2 | train |
| *Eragrostis nindensis* | PRJNA807505:<br>SRS12036931<br>SRS12036932 | PRJNA548367:<br>SRS4948778<br>SRS4948779<br>SRS4948781 | GCA_012490785.1 | train |

| | | | | |
|---|---|---|---|---|
| | | SRS4948782 | | |
| *Glycine max* | PRJNA657378:<br>SRS7209174<br>SRS7209175 | PRJNA753632:<br>SRR15458316<br>SRR15458321 | GCF_000004515.6 | train |
| *Malus domestica* | PRJNA821644:<br>SRS12449334<br>SRS12449335 | PRJNA267727:<br>SRS752518 | GCA_916612005.1 | train |
| *Marchantia polymorpha* | PRJNA597314:<br>SRR10879463<br>SRR10879464 | | GCA_003032435.1 | train |
| *Medicago truncatula* | PRJNA647765:<br>SRS7054112<br>SRS7054113<br>SRS7054114<br>SRS7054115<br>SRS7054116<br>SRS7054117<br>SRS7054118 | PRJNA783892:<br>SRS11159582<br>SRS11159583 | GCF_003473485.1 | val |
| *Oropetium thomaeum* | PRJNA807505:<br>SRS12036929<br>SRS12036933<br>SRS12036934<br>SRS12036935 | | GCA_001182835.1 | train |
| *Oryza brachyantha* | | PRJNA521886:<br>SRS4357813<br>SRS4357820<br>SRS4357828 | GCF_000231095.2 | train |
| *Oryza sativa* | PRJNA751145:<br>SRS9651698<br>SRS9651700<br>SRS9651701<br>SRS9651704<br>SRS9651708 | PRJNA386513:<br>SRS2419794<br>SRS2419800<br>SRS2419801 | GCF_001433935.1 | test |
| *Prunus persica* | | PRJNA381300:<br>SRS2712226<br>SRS2712231<br>PRJNA589110:<br>SRS5638125<br>SRS5638127<br>SRS5638129 | GCF_000346465.2 | train |
| *Pyrus x bretschneideri* | | PRJNA669907:<br>SRS7570511<br>SRS7570512<br>SRS7570517<br>SRS7570518<br>SRS7570519<br>SRS7570520<br>SRS7570521 | GCF_019419815.1 | train |
| *Sesamum indicum* | | PRJNA577518:<br>SRS5511999<br>SRS5512000<br>SRS5512001<br>SRS5512002 | GCF_000512975.1 | train |
| *Setaria italica* | | PRJNA391551:<br>SRS2307763<br>PRJNA486213:<br>SRS3675865 | GCF_000263155.2 | train |
| *Solanum lycopersicum* | PRJNA850391:<br>SRS13475373<br>PRJNA937410:<br>SRS16948284<br>SRS16948285<br>SRS16948288 | PRJNA624889:<br>SRS6475095<br>SRS6475096<br>SRS6475097 | GCF_000188115.5 | train |
| *Spirodela polyrhiza* | PRJNA527732:<br>SRS4500499<br>SRS4500501 | PRJNA527732:<br>SRS4500430 | GCA_900492545.1 | val |
| *Zea mays* | PRJNA697943: | PRJNA412230: | GCF_902167145.1 | train |

| | | |
|---|---|---|
| | SRS8775960 | SRR6077551 |
| | SRS8775993 | SRR6077553 |
| | SRS8775996 | SRR6077554 |

**Table S3: Tissues and treatments.** The tissues and/or treatments used for a given species is listed per NGS dataset and study (BioProject accession). Entirely unknown tissue and treatment is denoted with NaN.

| Species (scientific) | ATAC-seq (tissues/treatments) | ChIP-seq (tissues/treatments) |
|---|---|---|
| *Arabidopsis thaliana* | PRJNA394532:<br>Roots<br>PRJNA527732:<br>Leaves | PRJNA408288:<br>NaN<br>PRJNA535479:<br>Seedlings under cold treatment and following recovery |
| *Bigelowiella natans* | PRJNA753294:<br>Cell culture of unicellular algae | |
| *Brachypodium distachyon* | PRJNA661629:<br>Leaves under light and dark treatment | PRJNA661629:<br>Leaves under light and dark treatment |
| *Brassica napus* | PRJNA808238:<br>NaN | PRJNA687926:<br>Leaves |
| *Brassica oleracea* | | PRJNA687926:<br>Leaves |
| *Brassica rapa* | | PRJNA687926:<br>Leaves |
| *Chlamydomonas reinhardtii* | | PRJNA681680:<br>NaN |
| *Eragrostis nindensis* | PRJNA807505:<br>Desiccated leaves | PRJNA548367:<br>Leaves under drought and sufficient water treatment |
| *Glycine max* | PRJNA657378:<br>Leaves | PRJNA753632:<br>Leaves under normal and salt treatment |
| *Malus domestica* | PRJNA821644:<br>Unknown tissue under drought and sufficient water treatment | PRJNA267727:<br>Field-grown leaves |
| *Marchantia polymorpha* | PRJNA597314:<br>Thallus | |
| *Medicago truncatula* | PRJNA647765:<br>Roots at 0 h, 15 min, 30 min, 1 h, 2 h, 4 h, 8h after *Sinorhizobium meliloti* lipochitooligosaccharides treatment | PRJNA783892:<br>Whole aerial tissues harvested from 14–17-day-old plants at 4 h after dawn |
| *Oropetium thomaeum* | PRJNA807505:<br>Well-watered and desiccated leaves | |
| *Oryza brachyantha* | | PRJNA521886:<br>Aerial tissue |
| *Oryza sativa* | PRJNA751145:<br>Pistil and anther under low and normal temperature treatment | PRJNA386513:<br>Callus, leaves, and panicle |
| *Prunus persica* | | PRJNA381300:<br>Leaf and ripe fruit<br>PRJNA589110:<br>Vegetative bud |
| *Pyrus x bretschneideri* | | PRJNA669907:<br>Buds during dormancy transition |
| *Sesamum indicum* | | PRJNA577518:<br>Unknown tissue under light and dark treatment |
| *Setaria italica* | | PRJNA391551:<br>Leaf mesophyll<br>PRJNA486213:<br>Bundle sheath |
| *Solanum lycopersicum* | PRJNA850391:<br>4-weeks-old fourth leaves after 1 h of heat stress<br>PRJNA937410: | PRJNA624889:<br>Pericarp from the equatorial part of the fruit |

| | | |
|---|---|---|
| | Fruit | |
| *Spirodela polyrhiza* | PRJNA527732:<br>Leaves | PRJNA527732:<br>Leaves |
| *Zea mays* | PRJNA697943:<br>Unknown tissue (single cell) | PRJNA412230:<br>Root tips |

#### S3.2 Training parameters

**Table S4: Training sets.** For the first seven models only the species for which experimental ATAC-seq data of high quality was available were trained on. The same applies to the eighth model using ChIP-seq data. The combined model uses all available data.

| **Models** | U-Net<br>Hybrid<br>BiHybrid<br>BiHybrid_02<br>BiHybrid_03.1<br>BiHybrid_03.2 | BiHybrid_04 | BiHybrid_05 | Combined |
|---|---|---|---|---|
| **Comment** | Gap subsequences masked | Gap subsequences, unplaced scaffolds and non-nuclear sequences masked (see S1.6 Filtering flagged sequences) | see BiHybrid_04 | see BiHybrid_04 |
| **Training species** | *B. distachyon*<br>*B. napus*<br>*B. natans*<br>*E. nindensis*<br>*G. max*<br>*M. domestica*<br>*M. polymorpha*<br>*O. thomaeum*<br>*S. lycopersicum*<br>*Z. mays* | *B. distachyon*<br>*B. napus*<br>*B. natans*<br>*E. nindensis*<br>*G. max*<br>*M. domestica*<br>*M. polymorpha*<br>*O. thomaeum*<br>*S. lycopersicum*<br>*Z. mays* | *B. distachyon*<br>*B. napus*<br>*B. oleracea*<br>*B. rapa*<br>*C. reinhardtii*<br>*E. nindensis*<br>*G. max*<br>*M. domestica*<br>*O. brachyantha*<br>*P. bretschneideri*<br>*P. persica*<br>*S. indicum*<br>*S. italica*<br>*S. lycopersicum*<br>*Z. mays* | *B. distachyon*<br>*B. napus*<br>*B. natans*<br>*B. oleracea*<br>*B. rapa*<br>*C. reinhardtii*<br>*E. nindensis*<br>*G. max*<br>*M. domestica*<br>*M. polymorpha*<br>*O. brachyantha*<br>*O. thomaeum*<br>*P. bretschneideri*<br>*P. persica*<br>*S. indicum*<br>*S. italica*<br>*S. lycopersicum*<br>*Z. mays* |
| **Validation species** | *M. truncatula*<br>*S. polyrhiza* | *M. truncatula*<br>*S. polyrhiza* | *M. truncatula*<br>*S. polyrhiza* | *M. truncatula*<br>*S. polyrhiza* |

**Table S5: Model training parameters.** The parameter names are the exact naming convention used in Predmoter. The GitHub commit used is listed as Predmoter commit. A detailed explanation of the parameters can be found at: https://github.com/weberlab-hhu/Predmoter/blob/main/docs/Predmoter_options.md.

| **Para-meters** | **Model** | | | | | | | | |
|---|---|---|---|---|---|---|---|---|---|
| | U-Net | Hybrid | BiHybrid | BiHybrid_02 | BiHybrid_03.1 | BiHybrid_03.2 | BiHybrid_04 | BiHybrid_05 | Combined |
| Predmoter commit | cbc2256 | cbc2256 | cbc2256 | cbc2256 | cbc2256 | cbc2256 | c9ee6d7 | c9ee6d7 | c9ee6d7 |
| **Configu-ration para-meters** | | | | | | | | | |
| datasets | atacseq | atacseq | atacseq | atacseq | atacseq | atacseq | atacseq | h3k4me3 | atacseq, h3k4me3 |
| ram-efficient | true | true | true | true | true | true | true | true | true |
| **Model para-** | | | | | | | | | |

| meters | | | | | | | | | |
|---|---|---|---|---|---|---|---|---|---|
| model-type | cnn | hybrid | bi-hybrid | bi-hybrid | bi-hybrid | bi-hybrid | bi-hybrid | bi-hybrid | bi-hybrid |
| cnn-layers | 3 | 3 | 3 | 3 | 3 | 3 | 3 | 3 | 3 |
| filter-size | 64 | 64 | 64 | 64 | 64 | 64 | 64 | 64 | 64 |
| kernel-size | 18 | 18 | 18 | 18 | 18 | 18 | 18 | 18 | 18 |
| step | 3 | 3 | 3 | 3 | 3 | 3 | 3 | 3 | 3 |
| up | 2 | 2 | 2 | 2 | 2 | 2 | 2 | 2 | 2 |
| dilation | 1 | 1 | 1 | 1 | 1 | 1 | 1 | 1 | 1 |
| lstm-layers | / | 2 | 2 | 2 | 2 | 2 | 2 | 2 | 2 |
| hidden-size | / | 128 | 128 | 128 | 128 | 128 | 128 | 128 | 128 |
| bnorm | false | false | false | true | true | true | true | true | true |
| dropout | / | 0 | 0 | 0 | 0.3 | 0.5 | 0.3 | 0.3 | 0.3 |
| learning-rate | 0.001 | 0.001 | 0.001 | 0.001 | 0.001 | 0.001 | 0.001 | 0.001 | 0.001 |
| **Trainer/ callback para-meters** | | | | | | | | | |
| ckpt-quantity | avg_val_accuracy | avg_val_accuracy | avg_val_accuracy | avg_val_accuracy | avg_val_accuracy | avg_val_accuracy | avg_val_accuracy | avg_val_accuracy | avg_val_accuracy |
| save-top-k | -1 | -1 | -1 | -1 | -1 | -1 | -1 | -1 | -1 |
| stop-quantity | avg_train_loss | avg_train_loss | avg_train_loss | avg_train_loss | avg_train_loss | avg_train_loss | avg_train_loss | avg_train_loss | avg_train_loss |
| patience | 10 | 10 | 10 | 10 | 10 | 10 | 10 | 10 | 10 |
| batch size | 200 | 200 | 200 | 200 | 200 | 200 | 200 | 200 | 200 |
| device | gpu | gpu | gpu | gpu | gpu | gpu | gpu | gpu | gpu |
| num-devices | 2 | 2 | 2 | 2 | 2 | 2 | 2 | 2 | 2 |
| num-workers | 4 | 4 | 4 | 4 | 4 | 4 | 4 | 4 | 4 |

#### S3.3 Tabular results

**Table S6: Best models.** The best model was determined by the highest average validation Pearson correlation coefficient over all three replicates and all epochs (rounded to four decimal points). The epoch numbering uses Python convention, starting at zero. The listed models were used for testing. The models BiHybrid_03.1, BiHybrid_04, BiHybrid_05 and Combined were used for inference. The model checkpoint files can be found at: https://github.com/weberlab-hhu/predmoter_models.

| Model | Epoch | Seed | Validation Pearson's r |
|---|---|---|---|
| U-Net | 6 | 4227086911 | 0.4125 |
| Hybrid | 147 | 961333724 | 0.4370 |
| BiHybrid | 104 | 132709648 | 0.4884 |
| BiHybrid_02 | 36 | 132709648 | 0.5217 |
| BiHybrid_03.1 | 61 | 961333724 | 0.5336 |
| BiHybrid_03.2 | 53 | 961333724 | 0.5374 |
| BiHybrid_04 | 49 | 4227086911 | 0.5323 |
| BiHybrid_05 | 2 | 132709648 | 0.4387 |
| Combined | 20 | 4227086911 | 0.4835 |

**Table S7: Pearson's correlation for ATAC-seq predictions per species.** The predictions of the best models were compared with the experimental data per species. The resulting Pearson correlation coefficients were rounded to four decimal points. Gap subsequences were excluded from all test runs.

| Species (scientific) | Model | | | | | | | | | Split |
|---|---|---|---|---|---|---|---|---|---|---|
| | U-Net | Hybrid | BiHybrid | BiHybrid 02 | BiHybrid 03.1 | BiHybrid 03.2 | BiHybrid _03.1* | BiHybrid _04* | Com-bined* | |
| *Arabidopsis thaliana* | 0.2247 | 0.3338 | 0.4797 | 0.5916 | 0.6043 | 0.5881 | 0.6049 | 0.6122 | 0.6106 | test |
| *Bigelo-wiella natans* | 0.1043 | 0.3947 | 0.4876 | 0.5875 | 0.6182 | 0.5852 | 0.6182 | 0.6191 | 0.5608 | train |
| *Brachy-podium distachyon* | 0.5337 | 0.6203 | 0.6843 | 0.7265 | 0.7466 | 0.7371 | 0.7469 | 0.7404 | 0.7165 | train |
| *Brassica napus* | 0.2264 | 0.2778 | 0.3904 | 0.4403 | 0.4485 | 0.4371 | 0.4817 | 0.4855 | 0.4777 | train |

| | | | | | | | | | | |
|---|---|---|---|---|---|---|---|---|---|---|
| *Eragrostis nindensis* | 0.3114 | 0.4084 | 0.4650 | 0.5017 | 0.5323 | 0.5142 | 0.5323 | 0.5335 | 0.4930 | train |
| *Glycine max* | 0.5407 | 0.6222 | 0.6721 | 0.7124 | 0.7200 | 0.7138 | 0.7230 | 0.7245 | 0.7174 | train |
| *Malus domestica* | 0.2332 | 0.3524 | 0.4358 | 0.4829 | 0.5190 | 0.4917 | 0.5185 | 0.5262 | 0.4801 | train |
| *Marchantia polymorpha* | 0.3984 | 0.4671 | 0.5579 | 0.6037 | 0.6302 | 0.6121 | 0.6302 | 0.6334 | 0.5965 | train |
| *Medicago truncatula* | 0.4268 | 0.4498 | 0.4918 | 0.5363 | 0.5498 | 0.5497 | 0.5504 | 0.5489 | 0.5583 | val |
| *Oropetium thomaeum* | 0.4873 | 0.6482 | 0.7423 | 0.7713 | 0.7993 | 0.7816 | 0.7993 | 0.8022 | 0.7600 | train |
| *Oryza sativa* | 0.3743 | 0.4600 | 0.4818 | 0.5063 | 0.4926 | 0.4887 | 0.4930 | 0.4903 | 0.4472 | test |
| *Solanum lycoper-sicum* | 0.2897 | 0.3671 | 0.4361 | 0.4832 | 0.5037 | 0.4913 | 0.5077 | 0.5131 | 0.4964 | train |
| *Spirodela polyrhiza* | 0.3684 | 0.3970 | 0.4779 | 0.4766 | 0.4836 | 0.4994 | 0.4836 | 0.4813 | 0.4938 | val |
| *Zea mays* | 0.3185 | 0.4383 | 0.4854 | 0.5334 | 0.5469 | 0.5274 | 0.5496 | 0.5504 | 0.5400 | train |

* Flagged regions weren't tested.

**Table S8: Pearson's correlation for ChIP-seq predictions per species.** The predictions of the best models were compared with the experimental data per species. The resulting Pearson correlation coefficients were rounded to four decimal points. Gap subsequences and flagged regions were excluded from all test runs.

| **Species (scientific)** | **Model** | | **Split** |
|---|---|---|---|
| | BiHybrid_05 | Combined | |
| *Arabidopsis thaliana* | 0.7692 | 0.7641 | test |
| *Brachypodium distachyon* | 0.7391 | 0.7871 | train |
| *Brassica napus* | 0.6001 | 0.6222 | train |
| *Brassica oleracea* | 0.5602 | 0.5893 | train |
| *Brassica rapa* | 0.6328 | 0.6616 | train |
| *Chlamydomonas reinhardtii* | 0.7102 | 0.7988 | train |
| *Eragrostis nindensis* | 0.4914 | 0.5603 | train |
| *Glycine max* | 0.5097 | 0.5662 | train |
| *Malus domestica* | 0.4454 | 0.4919 | train |
| *Medicago truncatula* | 0.3957 | 0.3771 | val |
| *Oryza brachyantha* | 0.7991 | 0.8372 | train |
| *Oryza sativa* | 0.5918 | 0.6160 | test |
| *Prunus persica* | 0.6591 | 0.7055 | train |
| *Pyrus x bretschneideri* | 0.6088 | 0.6328 | train |
| *Sesamum indicum* | 0.7589 | 0.7895 | train |
| *Setaria italica* | 0.6559 | 0.7296 | train |
| *Solanum lycopersicum* | 0.3860 | 0.4227 | train |
| *Spirodela polyrhiza* | 0.5708 | 0.5699 | val |
| *Zea mays* | 0.2873 | 0.3294 | train |

#### S3.4 Peak statistics

**Table S9: Peak number statistics.** The number of peaks called from the mean experimental read coverage per test species and NGS dataset and of the predicted peaks per model, test species and NGS dataset are listed. The predicted and experimental peaks of both strands were summed.

| **Species (scientific)** | **ATAC-seq** | | | | **ChIP-seq** | | |
|---|---|---|---|---|---|---|---|
| | Experimental | BiHybrid_03.1 | BiHybrid_04 | Combined | Experimental | BiHybrid_05 | Combined |
| *A. thaliana* | 112,230 | 4,122 | 4,369 | 8,736 | 37,687 | 30,858 | 31,353 |
| *O. sativa* | 134,721 | 278,589 | 287,047 | 524,096 | 66,504 | 81,543 | 69,553 |

**Table S10: Peak length statistics.** The mean and median lengths of peaks called from the mean experimental read coverage per test species and NGS dataset and of the predicted peaks per model, test species and NGS dataset are listed. The lengths of the predicted and experimental peaks of both strands were used.

| **Species (scientific)** | **ATAC-seq** | **ChIP-seq** |
|---|---|---|

| | Experimental | BiHybrid_03.1 | BiHybrid_04 | Combined | Experimental | BiHybrid_05 | Combined |
|---|---|---|---|---|---|---|---|
| *A. thaliana* | 271/196 | 313/148 | 307/146 | 259/155 | 955/817 | 752/669 | 825/715 |
| *O. sativa* | 248/178 | 430/232 | 415/225 | 648/369 | 932/822 | 852/755 | 864/768 |

#### References

Buenrostro, Jason D., Giresi, Paul G., Zaba, Lisa C., Chang, Howard Y., and Greenleaf, William J., 'Transposition of Native Chromatin for Fast and Sensitive Epigenomic Profiling of Open Chromatin, DNA-Binding Proteins and Nucleosome Position', *Nature Methods*, 10/12 (2013), 1213–18

Hunter, John D., 'Matplotlib: A 2D Graphics Environment', *Computing in Science and Engineering*, 9/3 (2007), 90–95

Letunic, Ivica, and Bork, Peer, 'Interactive Tree Of Life (ITOL) v5: An Online Tool for Phylogenetic Tree Display and Annotation', *Nucleic Acids Research*, 49/W1 (2021), W293–96

Okonechnikov, Konstantin, Conesa, Ana, and García-Alcalde, Fernando, 'Qualimap 2: Advanced Multi-Sample Quality Control for High-Throughput Sequencing Data', *Bioinformatics*, 32/2 (2016), 292–94

Ramírez, Fidel, Ryan, Devon P., Grüning, Björn, Bhardwaj, Vivek, Kilpert, Fabian, Richter, Andreas S., et al., 'DeepTools2: A next Generation Web Server for Deep-Sequencing Data Analysis', *Nucleic Acids Research*, 44/W1 (2016), W160–65

Waskom, Michael L., 'Seaborn: Statistical Data Visualization', *Journal of Open Source Software*, 6/60 (2021), 3021

Zhang, Yong, Liu, Tao, Meyer, Clifford A., Eeckhoute, Jérôme, Johnson, David S., Bernstein, Bradley E., et al., 'Model-Based Analysis of ChIP-Seq (MACS)', *Genome Biology*, 9/9 (2008), 1–9
